# Astroglial Calcium Signaling Encodes Sleep Need in *Drosophila*

**DOI:** 10.1101/2020.07.04.187906

**Authors:** Ian D. Blum, Mehmet F. Keleş, El-Sayed Baz, Emily Han, Kristen Park, Skylar Luu, Habon Issa, Matt Brown, Margaret C.W. Ho, Masashi Tabuchi, Sha Liu, Mark N. Wu

## Abstract

Sleep is under homeostatic control, whereby increasing wakefulness generates sleep need and triggers sleep drive. However, the molecular and cellular pathways by which sleep need is encoded are poorly understood. In addition, the mechanisms underlying both how and when sleep need is transformed to sleep drive are unknown. Here, using *ex vivo* and *in vivo* imaging, we show in *Drosophila* that astroglial Ca^2+^ signaling increases with sleep need. We demonstrate that this signaling is dependent on a specific L-type Ca^2+^ channel and is required for homeostatic sleep rebound. Thermogenetically increasing Ca^2+^ in astrocytes induces persistent sleep behavior, and we exploit this phenotype to conduct a genetic screen for genes required for the homeostatic regulation of sleep. From this large-scale screen, we identify TyrRII, a monoaminergic receptor required in astrocytes for sleep homeostasis. TyrRII levels rise following sleep deprivation in a Ca^2+^-dependent manner, promoting further increases in astrocytic Ca^2+^ and resulting in a positive-feedback loop. These data suggest that TyrRII acts as a gate to enable the transformation of sleep need to sleep drive at the appropriate time. Moreover, our findings suggest that astrocytes then transmit this sleep need to the R5 sleep drive circuit, by upregulation and release of the interleukin-1 analog Spätzle. These findings define astroglial Ca^2+^ signaling mechanisms encoding sleep need and reveal dynamic properties of the sleep homeostatic control system.

## INTRODUCTION

The regulation of sleep by homeostatic forces is one of its defining features, and yet how sleep need is sensed and transduced into sleep drive remains poorly understood. The analysis of sleep homeostasis is guided by engineering control principles which posit that homeostatic systems comprise at least 3 components: a sensor that receives information about the state variable, an integrator that computes the difference between this state variable and a setpoint, and a downstream effector that responds to the integrator and directly manipulates the state variable[1]. Although the signal detected by “sleep sensors” remains debated, there is compelling evidence that a key stimulus for triggering sleep need is neuronal activity[2, 3]. For instance, studies in rodents and humans have demonstrated that tasks which activate specific regions in the brain will then locally promote an increase in the amplitude of electroencephalographic slow-wave activity, an established marker of sleep need[4–6].

More than a century ago, Cajal proposed that astrocytes, a subtype of glial cells, modulate neural connectivity across the sleep/wake cycle[7]. Since that time, emerging data have suggested that astrocytes play a key role in the regulation of sleep[8–11]. However, whether and how astrocytes sense and discharge sleep need is enigmatic. A number of special features of astrocytes make them well-suited for serving as “sensors” of sleep need. Astrocytes effectively tile the entire brain, and their processes form an intimate network around synapses and locally sample neural activity[12–15]. In addition, Ca^2+^ signaling plays an important role in astrocyte function, and modulation of intracellular Ca^2+^ levels is a broadly used mechanism for computing information[13, 16–20]. Finally, astrocytes appear to directly release neurotransmitters and other effector molecules (“gliotransmission”)[15, 21, 22] and, at least *in vitro*, have been shown to secrete sleep-promoting substances[23]. Thus, astrocytes could potentially sense sleep need-generating signals, perform relevant computations, and release signaling molecules to act on a downstream integrator circuit.

The morphology and functions of astrocytes are largely conserved in many animal species, including in the fruit fly *Drosophila melanogaster*[24]. In addition, sleep in *Drosophila* shares all defining behavioral criteria of sleep with mammals[25, 26]. Here, we use the fly model to characterize the role of astrocytes in sleep homeostasis and to delineate the mechanisms by which astroglial Ca^2+^ signals encode and transmit sleep need. Using *ex vivo* and *in vivo* imaging, we show that astrocytic Ca^2+^ levels vary with sleep need and demonstrate a critical role for this Ca^2+^ signaling in sleep homeostasis. Importantly, using a forward genetic approach, we identify Tyramine Receptor II (TyrRII), a monoaminergic receptor that is transcriptionally upregulated in astrocytes following sleep deprivation and which functions in a positive feedback Ca^2+^ loop to regulate sleep homeostasis. Our data further suggest that this astroglial Ca^2+^ signaling culminates in the upregulation of the secreted molecule Spätzle (Spz), the fly analog of mammalian interleukin 1 (IL-1); Spz then signals to a previously defined sleep drive circuit, to specifically control homeostatic rebound sleep. Together, these data support a model wherein astrocytes act as “sensors” of sleep need and encode sleep pressure under conditions of substantial sleep loss. Moreover, the underlying transcriptional and positive feedback signaling processes provide a mechanistic framework for conceptualizing the delay between and transformation of sleep need to sleep drive.

## RESULTS

### Ca^2+^ signaling in astrocytes correlates with sleep need and is necessary for sleep homeostasis

On balance, wakefulness is associated with greater neuronal activity[3, 6, 27–31]. Since astrocytes sense and respond to increases in neuronal activity through Ca^2+^ signaling mechanisms[10, 12, 15, 18, 22, 32, 33], we first asked whether Ca^2+^ levels in astrocytes vary according to sleep need. To do this, we expressed the genetically-encoded Ca^2+^ indicator CaMPARI2[34] in astrocytes and examined *ex vivo* CaMPARI signal in two different neuropil regions (superior medial protocerebrum, SMP, and antennal lobe, AL) at ZT0-3 (Zeitgeber time 0-3), ZT12-15 (mild increase in sleep need), and ZT0-3 following 12 hrs sleep deprivation (SD, strong increase in sleep need). Astrocytes exhibit distinct pools of intracellular Ca^2+^ (e.g., soma vs processes) that have different temporal kinetics and may participate in different signaling pathways[17, 18, 20, 33]. Thus, we analyzed CaMPARI signals in both of these locations. As shown in Figures 1A, 1B, and 1D, Ca^2+^ levels in the processes of astrocytes were elevated at ZT12-15, compared to ZT0-3, and further increased at ZT0-3 following SD. In the cell bodies, astrocytic Ca^2+^ levels were not elevated at ZT12-15 but were markedly elevated at ZT0-3 following SD (Figures 1A, 1C, and 1E). To address whether the increases in astrocyte Ca^2+^ concentrations simply reflected a stress response to mechanical SD, we also examined CaMPARI signal at ZT12-15 following 12 hrs of SD during the daytime. CaMPARI signals were not elevated under these conditions, compared to control flies at ZT12-15 (Figures 1B-1E). These data suggest that astrocyte intracellular Ca^2+^, particularly in the cell bodies, increases in a non-linear manner in response to time awake.

**Figure 1.**
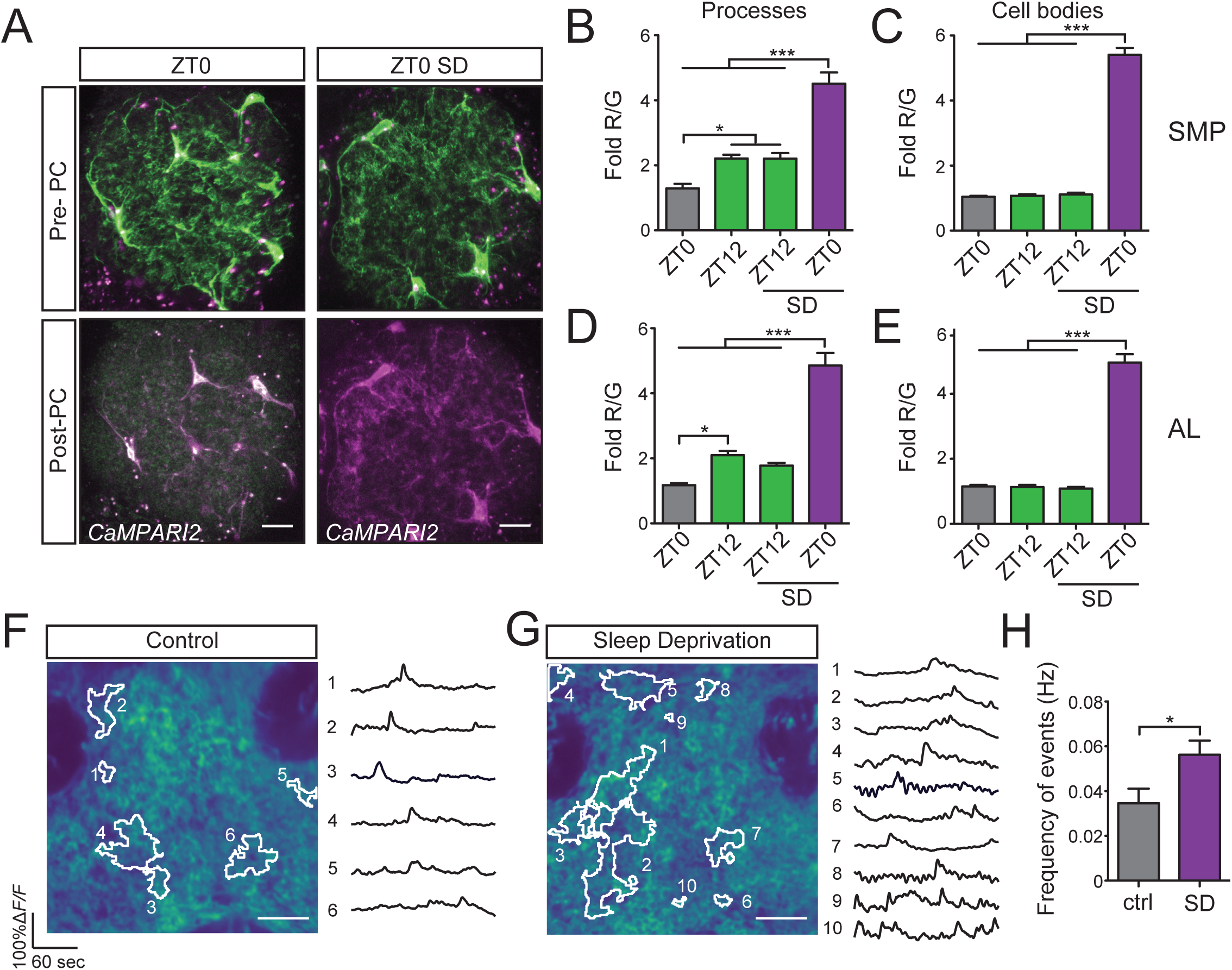
Ca^2+^ signaling in astrocytes correlates with sleep need. (A) Representative confocal images of pre-photoconversion (Pre-PC) and post-photoconversion (Post-PC) CaMPARI2 signal in the antennal lobe (AL) at ZT0 in the presence or absence of sleep deprivation (SD) from *R86E01-GAL4>UAS-CaMPARI2-L398T* flies. (B-E) CaMPARI2 signal (Fold R/G) from astrocyte processes (B and D) or cell bodies (C and E) from Superior Medial Protocerebrum (SMP) or AL from *R86E01-GAL4>UAS-CaMPARI2-L398T* flies at ZT0-3 or ZT12-15 under baseline conditions (AL: n=5 for ZT0-3 and n=6 for ZT12-15; SMP: n=5 for ZT0-3 and n=6 for ZT12-15) or after 12 hrs of SD (AL: n=6 for ZT0-3 and ZT12-15; SMP: n=6 for ZT0-3 and ZT12-15). (F and G) Representative 2-photon images (left) and event traces (right) of SMP from control (F) and sleep-deprived (G) *R86E01-GAL4>UAS-myr-GCaMP6s* flies at ZT0-2. Each image is the mean intensity of an entire recording from *in vivo* 2-photon Ca^2+^- imaging experiments. Data-driven regions of interest (ROIs) that were used to extract event traces are highlighted in white. Corresponding traces and ROIs are numbered. (H) Frequency of events for control (gray) and sleep-deprived (magenta) flies (n=9 flies for control and n=7 flies for SD) expressing membrane-bound GCaMP6s in astrocytes. Scale bars denote 10 μm in all images. For all bar graphs throughout manuscript, mean ± SEM is shown. In this and subsequent Figures, “*”, “**”, “***”, and “ns” denote *P*<0.05, *P*<0.01, *P*<0.001, and not significant, respectively.

We next examined whether changes in astrocytic Ca^2+^ following SD could be observed in living flies. We performed *in vivo* imaging of Ca^2+^ signals in astrocytic processes in the SMP region using myristoylated GCaMP (myr-GCaMP6s). The frequency of Ca^2+^ transients in the astrocytic processes was significantly increased following SD (Figures 1F-1H, S1A and S1B), whereas the event size and peak intensity of these Ca^2+^ transients were similar between these two conditions (Figures S1C and S1D). Taken together, our *ex vivo* and *in vivo* data argue that astrocytic Ca^2+^ signaling increases with greater sleep need.

To address whether astrocytic Ca^2+^ signaling is required for the homeostatic regulation of sleep, we sought to identify a molecule that fluxes Ca^2+^ and is important for this process. We conducted a candidate RNAi miniscreen of Ca^2+^-related channels, transporters, and exchangers and assayed for changes in sleep homeostasis. Knockdown of an L-type Ca^2+^ channel subunit (Ca-*α*1D) selectively in astrocytes led to a pronounced reduction in sleep rebound following SD (Figure 2A). These findings were next confirmed after backcrossing and with an additional RNAi line targeting Ca-*α*1D. Knockdown of Ca-*α*1D in astrocytes substantially reduced sleep recovery following SD (Figures 2B, 2C, and S2A) and led to a mild increase in baseline daily sleep time (Figure 2D) which was driven solely by an increase in nighttime sleep with no significant effects on sleep consolidation (Figures S2B-S2E). To address the functional role of Ca-*α*1D in regulating astrocytic Ca^2+^ levels, we measured CaMPARI signal in astrocytes in the AL following SD. As shown in Figures 2E-2G, knockdown of Ca-*α*1D suppressed the increased Ca^2+^ observed in astrocyte processes and cell bodies following SD. These data suggest that increases in astrocytic Ca^2+^ are required for normal homeostatic sleep rebound.

**Figure 2.**
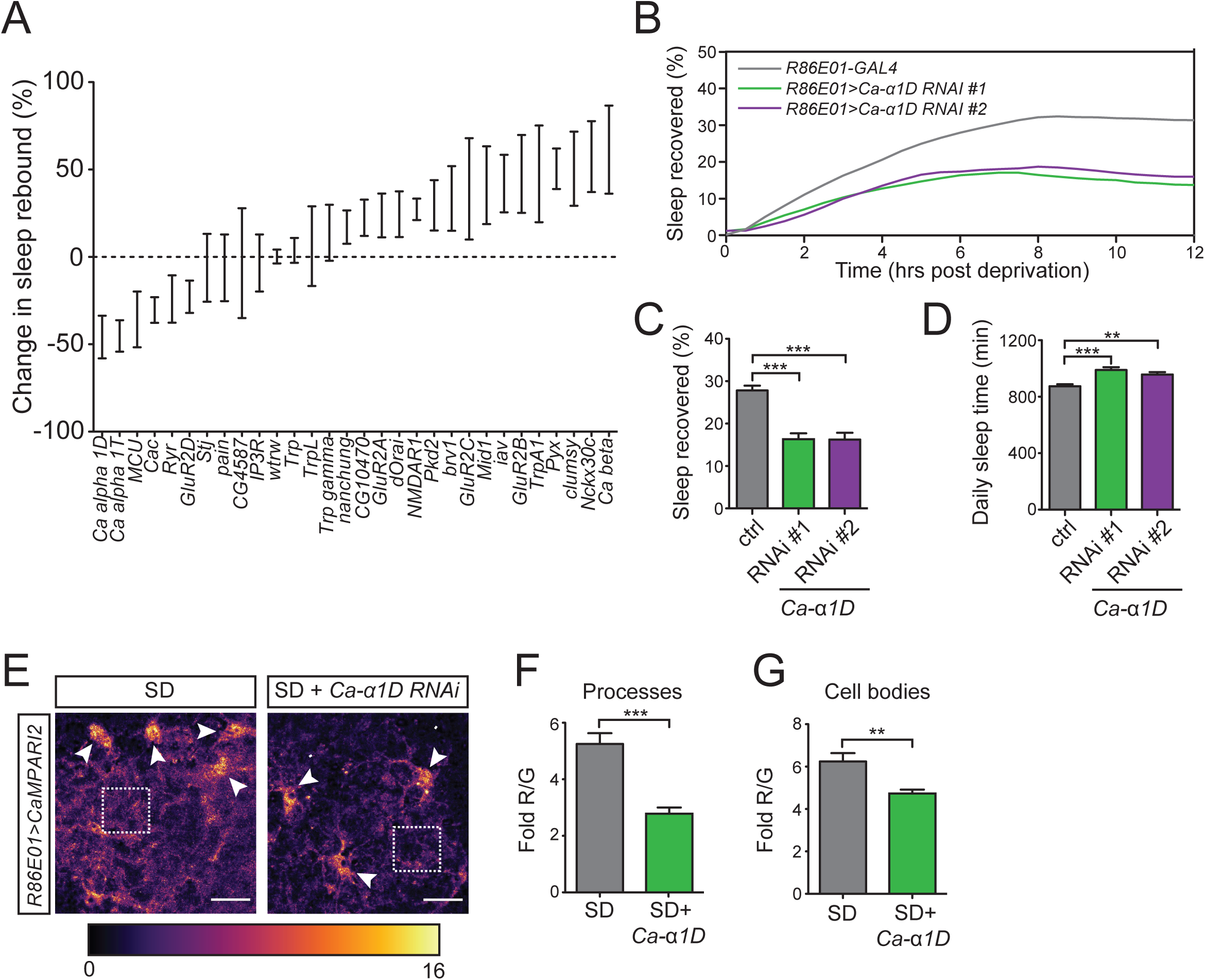
Ca^2+^ signaling in astrocytes is required for sleep homeostasis. (A) Mechanical deprivation miniscreen of astrocyte Ca^2+^-related channels, transporters, and exchangers. Relative change in sleep rebound for *R86E01-GAL4>UAS-RNAi* flies expressed as a percentage of sleep rebound observed in *R86E01-GAL4>UAS*-*empty vector* flies. (B) Sleep recovery curves for *R86E01-GAL4>ctrl* (gray) vs. *R86E01-GAL4>UAS-Ca-α1D-RNAi#1* (green) and *R86E01-GAL4>UAS-Ca-α1D-RNAi#2* (magenta) flies. (C and D) Sleep recovered (%) (C) and daily sleep amount (D) for *R86E01-GAL4>ctrl* (n=103), *R86E01-GAL4*>*UAS-Ca-α1D-RNAi#1* (n=53), and *R86E01-GAL4*>*UAS-Ca-α1D-RNAi#2* (n=35) flies. (E) Pixel-by-pixel heatmap of CaMPARI2 photoconversion signal in the AL region at ZT0-3 following 12 hr SD in *R86E01-GAL4>UAS-CaMPARI2-L398T* flies, in the presence or absence of *UAS-Ca-α1D-RNAi#1.* Dotted squares highlight Ca^2+^ signals from fine astrocyte processes, and white arrows denote cell bodies. Scale bars denote 10 μm. (F and G) Quantification of average CaMPARI2 signal (Fold R/G) from ROIs targeting astrocyte processes (F) or cell bodies (G) represented in (E).

### Ectopic astroglial Ca^2+^ signaling triggers both acute and delayed increases in sleep behavior

To examine the consequences of increasing Ca^2+^ levels within astrocytes on sleep behavior, we expressed the temperature-sensitive cation channel dTrpA1 in astrocytes. Interestingly, we found that elevating Ca^2+^ in astrocytes (*alrm-GAL4*) during the night led to two phenotypes: a rapid increase in nighttime sleep during the heat pulse and a persistent increase in daytime sleep following the cessation of the heat pulse. These two phenotypes were also observed using an additional astrocyte driver (*R86E01-GAL4*) and were reminiscent of the distinct phenotypes observed with activation of the ExFl2 sleep effector circuit (*R72G06-GAL4*) and the R5 (previously termed R2) sleep drive circuit (*R58H05-AD;R46C03-DBD*), respectively (Figures 3A-3C)[35, 36]. We manually scored fly behavior using video to demonstrate that flies were truly inactive (and not grooming or feeding) during and after thermogenetic activation. These data demonstrated that immobility was specifically increased by these manipulations (Figure S3A). To further confirm that the immobility measured reflects sleep and not simply immobility or paralysis, we assessed arousal threshold. Both during and after thermogenetic activation of astrocytes, flies demonstrated an increased arousal threshold to mild and moderate stimuli but were fully responsive to strong stimuli (Figure S3B). Finally, to ensure that the sleep induced by thermogenetic activation of astrocytes is reversible, we quantified sleep for several days after heat treatment (Figures S3C-S3E). In general, sleep normalizes by the night directly following the “sleep rebound,” with the exception of a decrease of daytime sleep the following day (“Day 2”), which may represent “negative sleep rebound”, a phenomena previously observed in both mammals and flies[37–39].

**Figure 3.**
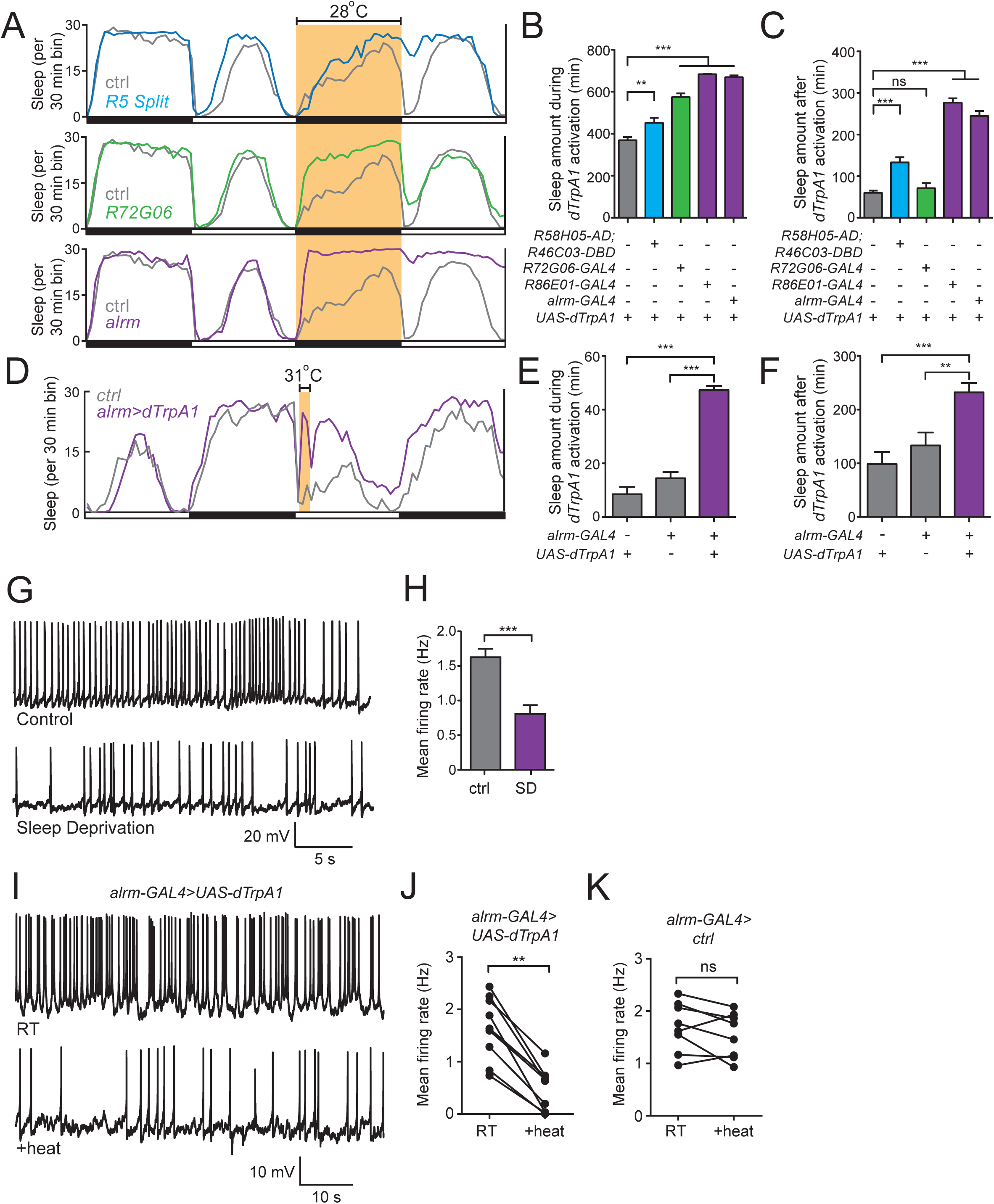
Astrocyte activation induces both proximate and delayed sleep and inhibits neural activity. (A) Sleep profile of *ctrl>UAS-dTrpA1* flies (gray) vs *R58H05-AD;R46C03-DBD>UAS-dTrpA1* (blue, upper panel), *R72G06-GAL4>UAS-dTrpA1* (green, middle panel) or *alrm-GAL4>UAS-dTrpA1* flies (magenta, lower panel). Highlighted period denotes 12 hr dTrpA1 activation at 28°C. Note that the controls in the 3 sleep profiles are from the same dataset. (B and C) Sleep amount over a 12 hr period during (B) or 6 hr after (C) dTrpA1 activation for *ctrl>UAS-dTrpA1* (n=81), *R58H05-AD;R46C03-DBD>UAS-dTrpA1* (n=17), *R72G06-GAL4>UAS-dTrpA1* (n=51), *R86E01-GAL4>UAS-dTrpA1* (n=29), and *alrm-GAL4>UAS-dTrpA1* (n=40). (D) Sleep profile of *ctrl>UAS-dTrpA1* (gray) and *alrm-GAL4>UAS-dTrpA1* (magenta). Highlighted period denotes 1 hr dTrpA1 activation at 31°C. (E and F) Sleep amount during (E) and after (F) 1 hr dTrpA1 activation for *ctrl>UAS-dTrpA1* (n=32), *alrm-GAL4*>*ctrl* (n=32), and *alrm-GAL4>UAS-dTrpA1* (n=28) flies. (G and H) Representative whole-cell recording traces (G) and mean firing rate (H) of spontaneous action potentials from l-LNvs in *PDF-GAL4>UAS-CD8::GFP* flies from ZT0-2 under baseline (“ctrl”, n=8) or 12 hr sleep-deprived (“SD”, n=8) conditions. (I-K) Representative whole-cell recording traces (I) and mean firing rate of spontaneous action potentials of l-LNvs in *alrm-GAL4>UAS-dTrpA1;PDF-LexA>LexAop2-CD2::GFP* (n=9) (J) or *alrm-GAL4>ctrl, PDF-LexA*>*LexAop2-CD2::GFP* (n=8) (K) flies from ZT0-2 at 22°C (“RT”) and 29°C (“+heat”).

We next asked whether this persistent “sleep rebound” could be induced by a shorter period of astrocyte activation, similar to our previous findings with the R5 sleep drive circuit[36]. Indeed, 1 hr heat activation of dTrpA1 in astrocytes also triggered a persistent increase in sleep, suggestive of an increase in sleep drive. Interestingly, however, unlike the case for R5 neurons where a monophasic increase in sleep was induced following the heat treatment[36], 1 hr activation of astrocytes led to a biphasic sleep response—a rapid increase in sleep during the heat treatment followed by a delayed increase in sleep after the heat treatment (Figures 3D-3F). Similar observations were made using *R86E01-GAL4* (Figures S3F and S3G). These data suggest that activation of astrocytes is sufficient for inducing sleep behavior and generating homeostatic sleep drive.

Astrocytes release various neurotransmitters and signaling molecules to modulate synaptic function[14, 15, 21, 22]. What effect might Ca^2+^ signaling in astrocytes have on neural activity? To address this, we performed patch-clamp recordings of the large ventrolateral clock neurons (l-LNvs). As clock neurons that modulate arousal[40–42], the firing rate of the l-LNvs is under circadian control[43–45], but their sensitivity to sleep need in unknown. Therefore, we first examined whether l-LNv firing is altered following 12 hrs of SD; indeed, their firing rate at ZT0-2 was significantly reduced and their resting membrane potential was hyperpolarized after SD (Figures 3G, 3H, and S4A). We then assessed the impact of dTrpA1 activation of astrocytes on l-LNv activity. *alrm>dTrpA1*-mediated astrocyte activation led to a significant reduction in spiking frequency and resting membrane potential, whereas heat treatment alone (*UAS-dTrpA1*) had no effect on firing rate (Figures 3I-3K, S4B, and S4C). Similar observations were made using *R86E01-GAL4* (Figures S4D and S4E). These data suggest that activation of astrocytes can result in inhibition of arousal-promoting systems.

### TyrRII is required for astroglial control of sleep homeostasis

Little is known about the molecular pathways by which astrocytes regulate sleep. The use of forward genetic screens to identify novel genes critical for sleep homeostasis has been hindered by the difficulty of performing sleep deprivation robustly and reproducibly on a large-scale. To circumvent this problem and identify astrocytic genes required for sleep homeostasis, we capitalized on our finding of persistent sleep drive following astrocyte activation and performed a screen for genes required for this phenotype. From a screen of ∼3,200 RNAi lines, we identified ∼40 lines that reproducibly suppressed this sleep phenotype (Figure 4A). Interestingly, we found that knockdown of the largely uncharacterized receptor TyrRII markedly suppressed the persistent sleep phenotype seen with activation of astrocytes (Figures 4A-4C). To address whether TyrRII expression in astrocytes is directly required for the homeostatic regulation of sleep, we assessed sleep recovery following 12 hrs of mechanical SD. Sleep recovery, but not baseline daily sleep, was significantly reduced when TyrRII was knocked down in astrocytes, compared to controls (Figures 4D-4F and S5A). In contrast, knockdown of TyrRII in astrocytes did not yield consistent effects on baseline daytime sleep, nighttime sleep, or sleep consolidation (Figures S5B-S5E).

**Figure 4.**
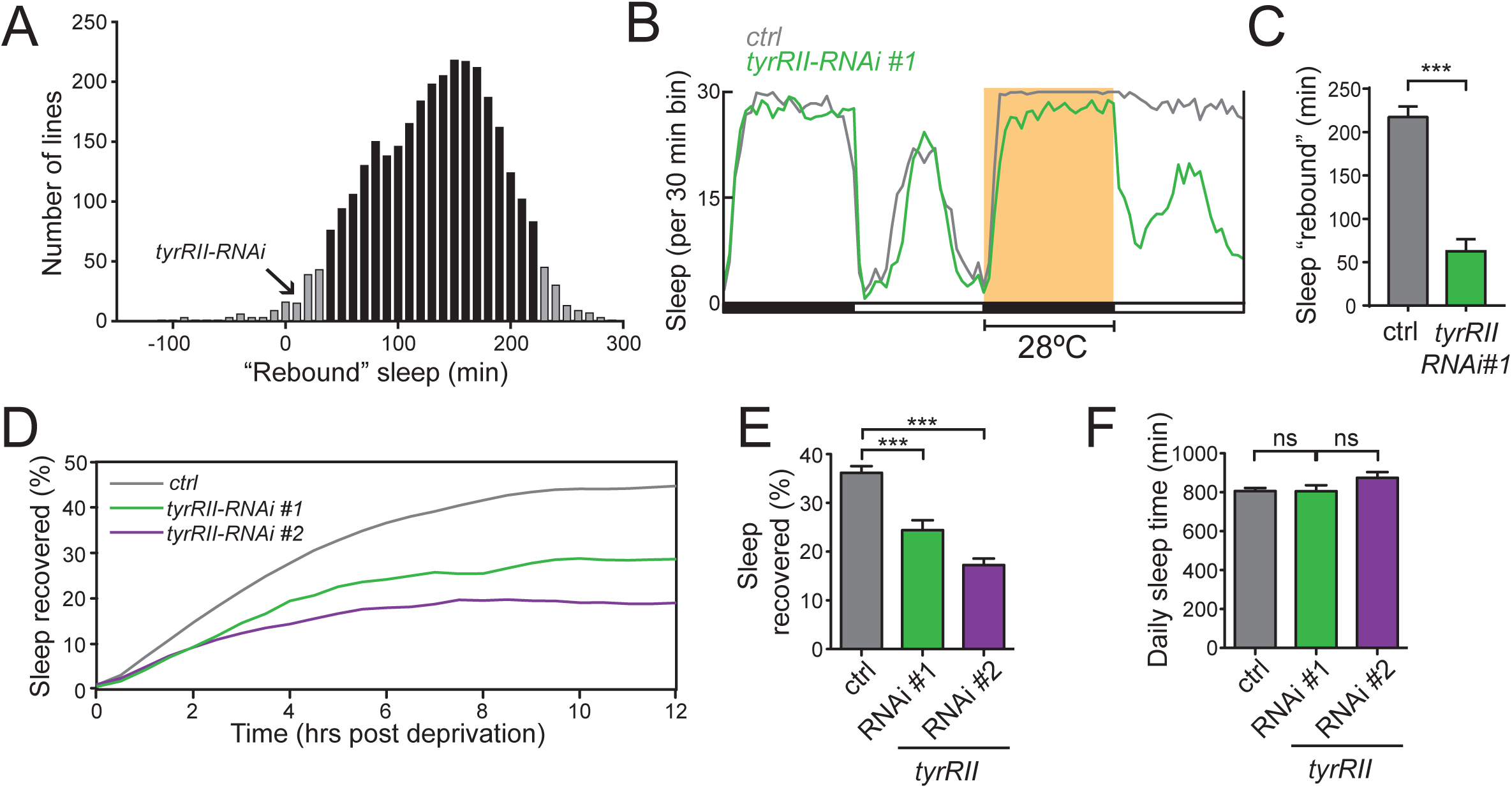
TyrRII is required in astrocytes for homeostatic sleep rebound. (A) Histogram for RNAi genetic screen for *alrm-GAL4>UAS-dTrpA1, UAS-RNAi* flies (n=3,150 genes, n=4 flies per genotype) showing “rebound” sleep (6 hr post-activation) induced by 1 hr of heat (31°C) from ZT0-1. The amount of “rebound sleep” for *alrm-GAL4>UAS-TyrRII-RNAi#1-*expressing flies is noted. Bars in gray denote values lying +/- 2.5 SD from the mean. (B) Sleep profile of *alrm-GAL4>UAS-dTrpA1, ctrl* (gray), and *alrm-GAL4>UAS-dTrpA1, UAS-TyrRII-RNAi#1* (green) flies. Highlighted period denotes 12 hr dTrpA1 activation at 28°C. (C) Sleep amount for the 6 hr period after dTrpA1 activation for *alrm-GAL4>UAS-dTrpA1, ctrl* (n=19), and *alrm-GAL4>UAS-dTrpA1*, *UAS-TyrRII-RNAi#1* (n=24). (D) Sleep recovery curves for *R86E01-GAL4>ctrl* (gray) vs *R86E01-GAL4>UAS-TyrRII-RNAi#1* (green) and *R86E01-GAL4>UAS-TyrRII-RNAi#2* (magenta) flies after overnight (12 hr) SD. (E and F) Sleep recovered (%) (E) and daily sleep amount (F) for *R86E01-GAL4>ctrl* (n=81) and *R86E01-GAL4>UAS-TyrRII-RNAi* #1 (n=33), and *R86E01-GAL4>UAS-TyrRII-RNAi* #2 (n=36).

### TyrRII is upregulated with sleep loss and participates in a positive feedback Ca^2+^- signaling mechanism

TyrRII has previously been shown to be broadly responsive to a variety of monoamines and, given that monoamine release is generally associated with arousal[46–49], this system could represent an elegant mechanism for astrocytes to track wakefulness and consequently homeostatic sleep need. Because monoaminergic receptor expression is often tightly regulated by diverse signaling mechanisms[50], we examined whether *tyrRII* expression varied according to sleep need. We assessed astrocyte expression of *tyrRII* mRNA using TRAP-qPCR (Translating Ribosome Affinity Purification followed by Quantitative PCR)[51], which we performed by expressing eGFP-RpL10a in astrocytes and immunopurifying actively translating mRNA using magnetic beads coated with anti-GFP antibodies (Figure 5A). As expected, immunoprecipitates were dramatically enriched for a glial marker (*repo*), compared to a neuronal marker (*nSyb*) (Figure 5B). Interestingly, *tyrRII* mRNA was markedly elevated ∼ 10-fold after sleep deprivation (Figure 5C). This increase could be recapitulated by dTrpA1 activation of astrocytes, suggesting that the sleep need-dependent increase in *tyrRII* is Ca^2+^-dependent (Figures 5D and 5E).

**Figure 5.**
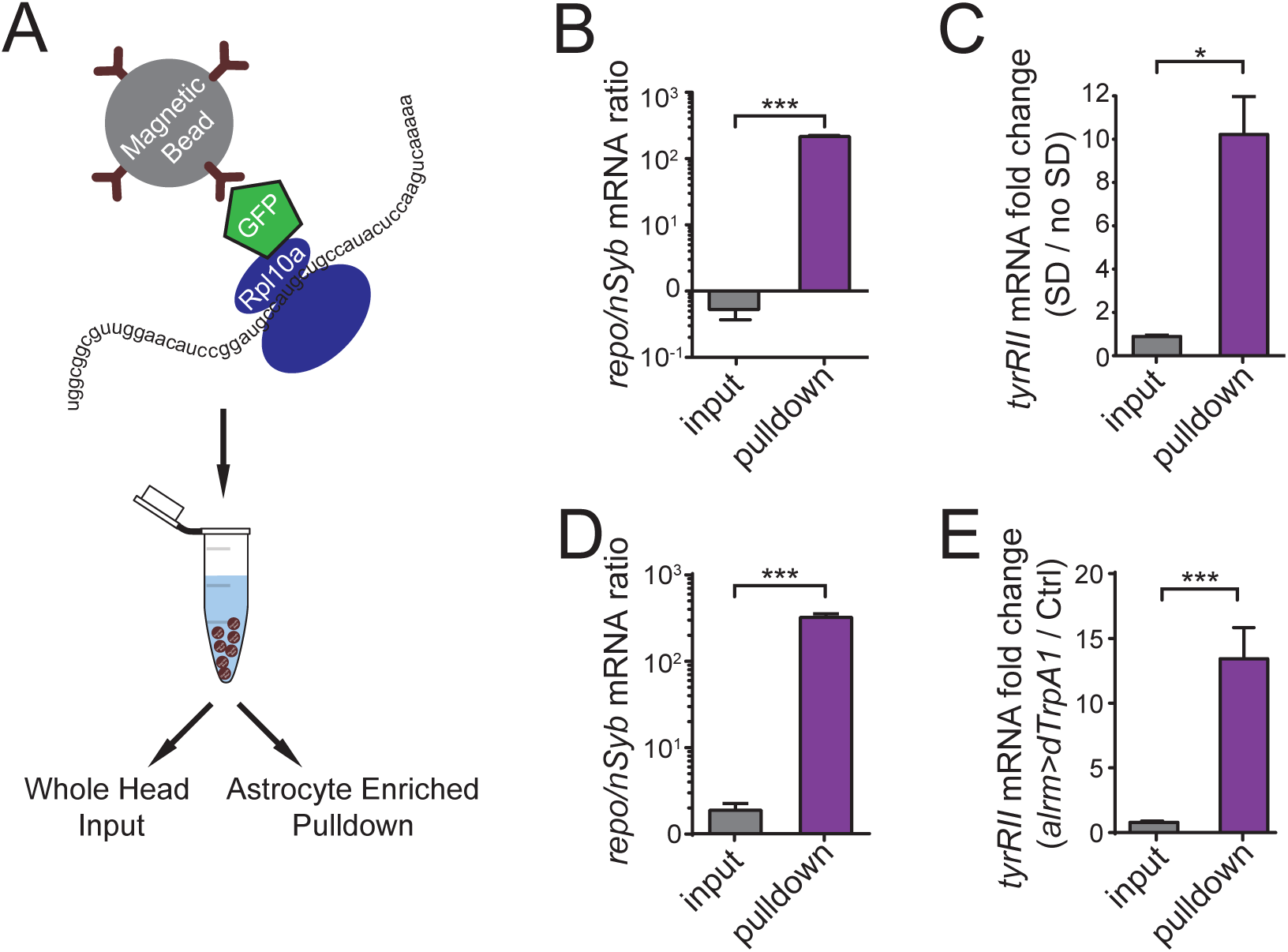
*tyrRII* mRNA is upregulated with sleep deprivation and astrocyte activation. (A) Schematic of translating ribosomal affinity purification (TRAP) procedure for isolating actively translating mRNA from genetically-defined astrocytes in whole fly heads. (B) Relative ratio of astrocyte marker (*repo*) and neural marker (*nSyb*) mRNA level in whole head flowthrough (input, n=3 replicates) vs astrocyte-TRAP (pulldown, n=3 replicates) samples from sleep deprivation experiment. (C) Relative change in *tyrRII* mRNA level in sleep-deprived (“SD”, n=3 replicates) vs non-sleep deprived (“no SD”, n=3 replicates) flies from astrocyte-TRAP (pulldown) vs whole head (input) samples. (D) Relative ratio of astrocyte marker (*repo*) and neural marker (*nSyb*) mRNA level in whole head flowthrough (input, n=3 replicates) vs astrocyte-TRAP (pulldown, n=3 replicates) samples from experiments where astrocytes are thermogenetically activated. (E) Relative fold change in *tyrRII* mRNA level in thermogenetically activated *alrm-GAL4>UAS-dTrpA1* (n=3 replicates) vs *ctrl>UAS-dTrpA1* (“ctrl”, n=3 replicates) flies from astrocyte-TRAP (pulldown) vs whole head (input) samples.

We next asked whether TyrRII protein levels were also increased following sleep deprivation. To address this question, we used a MiMIC transposon insertion[52], where GFP is fused in-frame into the 2^nd^ extracellular loop of the TyrRII protein. First, we assayed baseline and rebound sleep in homozygote *Mi{PT-GFSTF.2}TyrRII^MI12699^* flies and found no differences compared to heterozygous controls, suggesting that the GFP insertion does not disrupt protein function or localization (Figures S5F-S5H). Consistent with our findings with *tyrRII* transcript, TyrRII::GFP expression was substantially increased following 12 hr SD vs non-SD controls, both in terms of number and size of puncta (Figures S5I-S5K). Interestingly, super-resolution microscopy revealed that the sleep deprivation-mediated increase in TyrRII-GFP expression within the AL neuropil was largely localized to bulbous structures within astrocytic processes (Figure S5L). These structures were observed by *R86E01-GAL4*-driven tdTomato expression independent of sleep need-state, suggesting that they are not a direct result of either sleep deprivation or the expression of TyrRII::GFP.

Our previous data suggested that the upregulation of *tyrRII* transcript following SD is dependent on astrocytic intracellular Ca^2+^ (Figure 5E). Moreover, thermogenetic activation of astrocytes resulted in a marked increase in TyrRII::GFP puncta number and a significant, but less pronounced, increase in puncta size (Figures 6A-6C). To directly test whether astrocytic Ca^2+^ signaling is required for the sleep need-induced elevation in TyrRII::GFP, we performed RNAi knockdown of Ca-*α*1D in astrocytes and quantified TyrRII::GFP expression before and after SD. As shown in Figures 6D-6F, TyrRII::GFP expression following SD, but not under baseline conditions, was significantly reduced with concomitant knockdown of Ca-*α*1D, compared to controls. Prior studies have shown that monoaminergic signaling induces Ca^2+^ elevations in astrocytes in both flies and mice[53, 54]. Thus, TyrRII might produce an amplifying positive-feedback loop in astrocytes-- not only does TyrRII expression depend on intracellular Ca^2+^ levels, but the higher levels of TyrRII expression following SD contribute to further increases in Ca^2+^ levels. To address this possibility, we examined whether loss of TyrRII suppressed the elevation of intracellular Ca^2+^ seen in astrocytes following SD. Following SD, CaMPARI signal in both astrocyte processes and cell bodies was substantially reduced in *R86E01-GAL4>UAS-TyrRII-RNAi* animals, compared to controls (Figures 6G-6I). Taken together, these data suggest that, as sleep need accrues during protracted arousal, TyrRII amplifies Ca^2+^ signaling in a positive-feedback manner, priming astrocytes and sensitizing them to monoamines.

**Figure 6.**
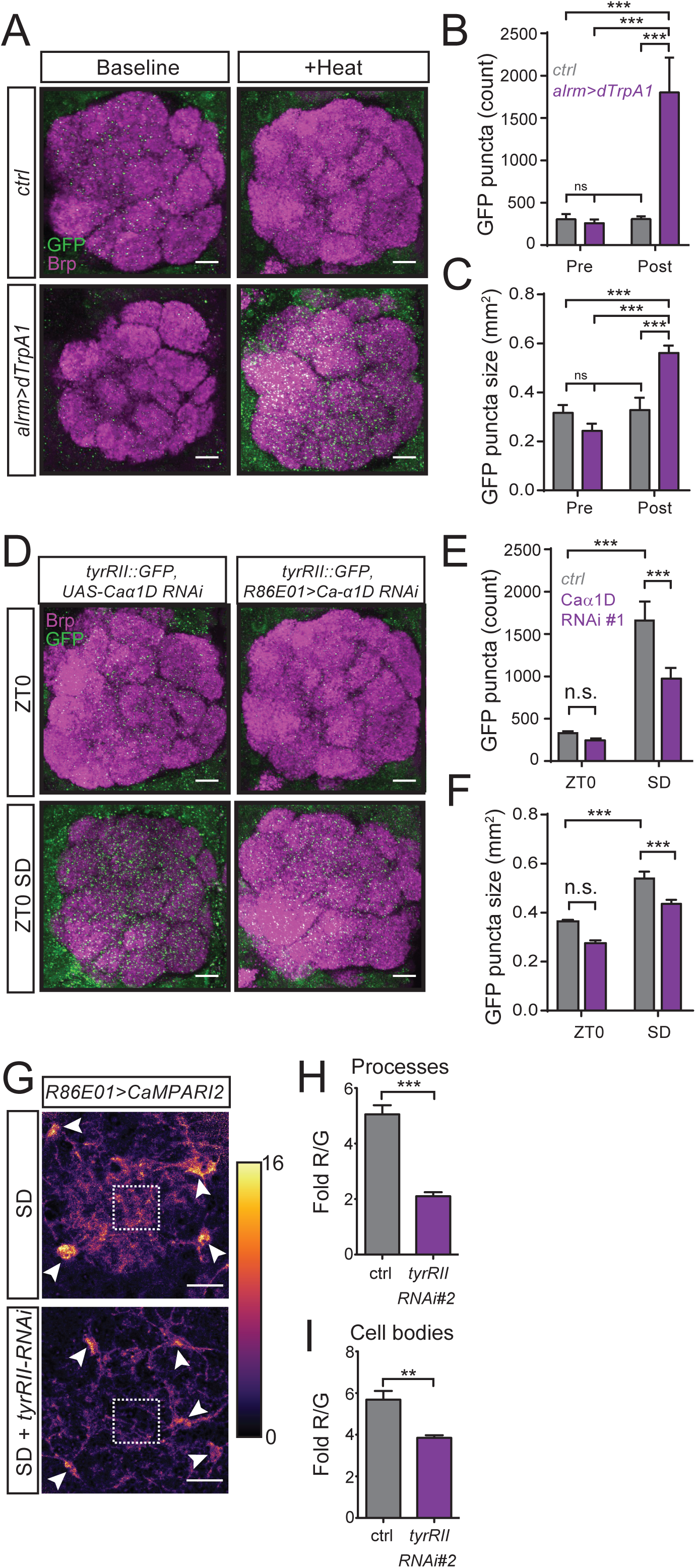
Astrocytic TyrRII is upregulated with sleep need and participates in a positive-feedback calcium signaling mechanism. (A) Representative images of TyrRII::GFP signal at the AL from *alrm>dTrpA1, Mi{PT-GFSTF.2}TyrRII^MI12699^/+* or *ctrl>UAS-dTrpA1, Mi{PT-GFSTF.2}TyrRII^MI12699^*/+ flies. Whole-mount brains were immunostained with anti-GFP (green) and anti-BRP (magenta) at ZT0-1 in the presence (*alrm>dTrpA1*) or absence (*ctrl>dTrpA1*) of thermogenetic activation (“+Heat”, 12 hrs at 28°C). (B and C) Number (B) and size (C) of TyrRII::GFP puncta in the AL at ZT1 in the presence (*alrm>dTrpA1*) or absence (*ctrl>dTrpA1*) of before (“Pre”) and after (“Post”) thermogenetic activation (n=5 for all groups and conditions). (D) Representative images of TyrRII::GFP signal at the AL following 12 hr sleep deprivation from *ctrl>UAS-Ca-α1D RNAi#1, Mi{PT-GFSTF.2}TyrRII^MI12699^/+* or *R86E01-GAL4>UAS-Ca-α1D RNAi#1, Mi{PT-GFSTF.2}TyrRII^MI12699^/+* flies. Whole-mount brains were collected from ZT0-1 and immunostained with anti-GFP (green) and anti-BRP (magenta). (E and F) Number (E) and size (F) of TyrRII::GFP puncta in the AL under baseline (ZT0) or SD conditions with (magenta) or without (gray) *Ca-α1D* knockdown (n=8 for all groups and conditions). (G) Pixel-by-pixel heatmap of CaMPARI2 photoconversion signal in the AL region from ZT0-3 following 12 hr SD in *R86E01-GAL4>UAS-CaMPARI2-L398T* flies, in the presence (n=9) and absence (n=10) of *UAS-TyrRII-RNAi#2*. (H and I) Quantification of CaMPARI2 signal (Fold R/G) from astrocyte processes (H) or cell bodies (I). Scale bars denote 20 μm in (A) and (D) and 10 μm in (G).

### Astrocyte-derived Spätzle transmits sleep need to the R5 sleep drive circuit

Not only are the underlying molecular pathways unclear, but the circuit mechanisms by which astrocytes signal sleep need are also unknown. In principle, such signaling could occur locally to neurons throughout the brain. Alternatively, sleep need could be integrated and transmitted to a central neural circuit. We first asked whether a specific signaling molecule is released from astrocytes to convey sleep need. *spätzle* (*spz*), the *Drosophila* analog of IL-1, was a “hit” in our dTrpA1-mediated RNAi screen, and, because IL-1 has previously been implicated in the homeostatic regulation of sleep in mammals[55], we focused on this gene. We found that knockdown of *spz* in astrocytes significantly reduced sleep recovery following SD (Figures 7A, 7B, and S6A), while not affecting daily sleep time, daytime sleep, nighttime sleep, or sleep consolidation under baseline conditions (Figure 7C and S6B-S6E). We next investigated whether *spz* expression was altered in response to changes in sleep need. We again performed TRAP-qPCR and found that *spz* transcript was increased ∼9-fold in astrocytes following SD (Figure 7D) or thermogenetic activation (Figure S6F), supportive of a role for astrocytic Spz in relaying sleep need.

**Figure 7.**
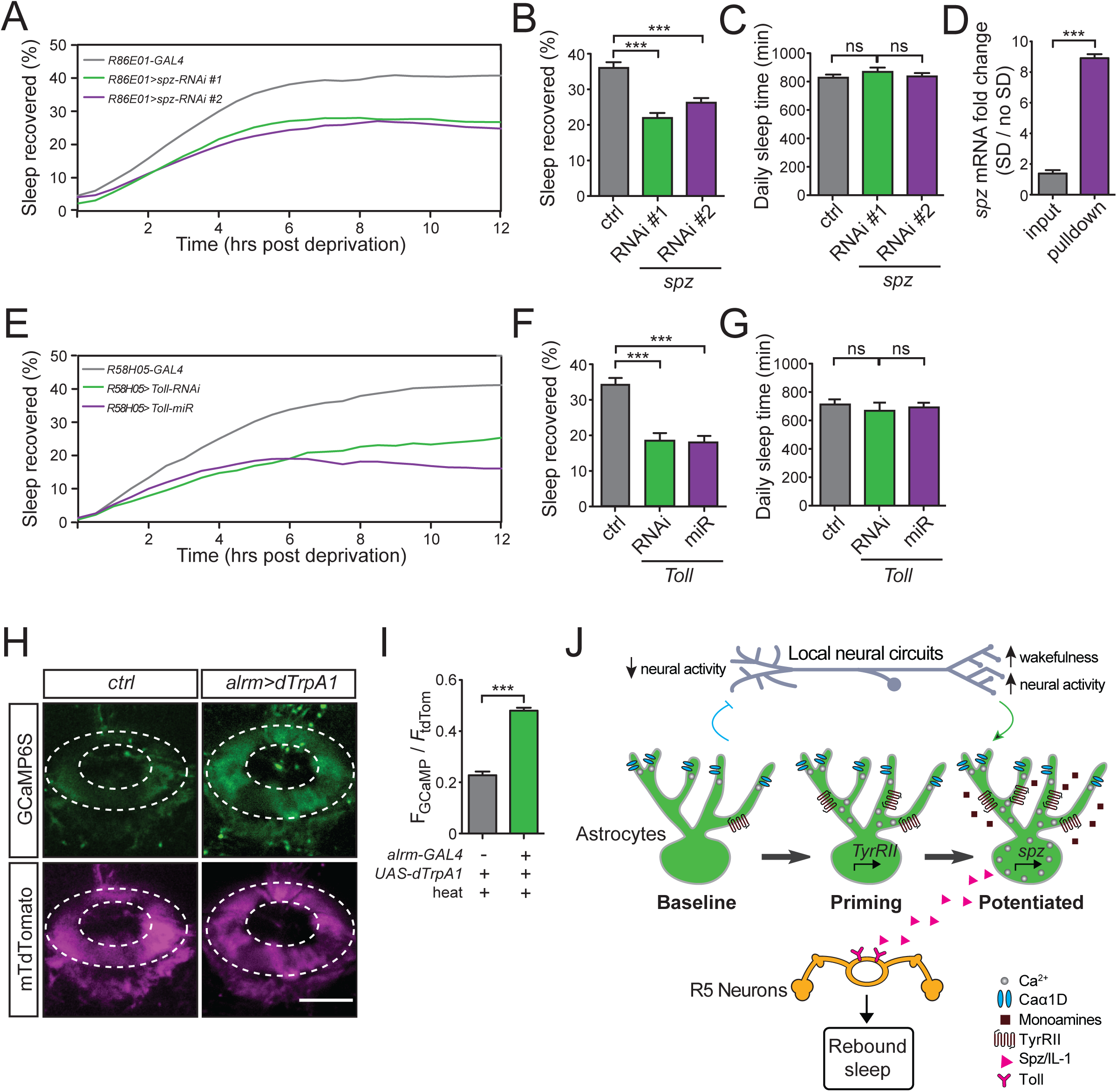
Astrocyte-derived Spätzle, an IL-1 analog, signals sleep need to the R5 sleep drive circuit via the Toll receptor. (A) Sleep recovery curve for *R86E01-GAL4>ctrl* (gray), *R86E01-GAL4>UAS-spz-RNAi#1* (green), and *R86E01-GAL4>UAS-spz-RNAi#2* (magenta) flies. (B and C) Sleep recovered (%) (B) and daily sleep amount (C) for *R86E01-GAL4>ctrl* (n=46), *R86E01-GAL4*>*UAS-spz-RNAi#1* (n=40), and *R86E01-GAL4*>*UAS-spz-RNAi#2* (n=56) flies. (D) Relative change in *spz* mRNA level in sleep-deprived (“SD”, n=3 replicates) vs non-sleep deprived (“no SD”, n=3 replicates) flies from astrocyte-TRAP (pulldown) vs whole head (input) samples. (E) Sleep recovery curve of *R58H058-GAL4>ctrl* (gray), *R58H05-GAL4>UAS-Toll-RNAi* (green), and *R58H05-GAL4>UAS-Toll-miR* (magenta). (F and G) Sleep recovered (%) (F) and daily sleep amount (G) for *R58H05-GAL4>ctrl* (n=103), *R58H05-GAL4>UAS-Toll-RNAi* (n=51), and *R58H05-GAL4>UAS-Toll-miR* (n=56) flies. (H and I) Representative images of GCaMP (upper panels) and tdTomato (lower panels) fluorescence intensity (H) and relative GCaMP fluorescence intensity (I) in the R5 ring of *ctrl>QUAS-dTrpA1; R58H05-GAL4>UAS-GCaMP6s, UAS-CD4::tdTomato* (ctrl, n=5) vs *alrm-QF2>QUAS-dTrpA1; R58H05-GAL4>UAS-GCaMP6s, UAS-CD4::tdTomato* (n=4) flies at ZT3-4 after 12 hrs of heat treatment from ZT12-24 at 28°C. For (H), dashed lines indicate the R5 ring, and scale bar denotes 20 μm. (J) Model for astroglial Ca^2+^ signaling in the homeostatic regulation of sleep. Neural activity during wakefulness is sensed by local astrocytes, resulting in increased Ca^2+^ in the processes, which requires specific voltage-gated Ca^2+^ channels (“Baseline”). Sleep loss generates protracted calcium signaling and leads to upregulation of TyrRII, sensitizing astrocytes to the actions of monoamines that are associated with wakefulness and further increasing Ca^2+^ levels in these cells (“Priming”). When sufficient sleep need has accumulated, as measured by heightened levels of astroglial Ca^2+^, transcription/translation of Spz is upregulated (“Potentiated”). Spz is then released and acts on Toll receptors in the R5 neurons to trigger global sleep drive.

We previously identified a neural circuit, comprising R5 ellipsoid body (EB) ring neurons, that encodes sleep drive[36]; we hypothesized that astrocytes may convey information regarding sleep need to these neurons. Moreover, recent data suggest that Toll, the receptor for Spz, is enriched within the EB in the adult brain[56]. Knockdown of Toll in R5 neurons led to a significant reduction in sleep recovery following SD (Figures 7E, 7F, and S6G), but not daily sleep time, or daytime sleep, nighttime sleep, and sleep consolidation under baseline conditions (Figures 7G and S6H-S6K). We previously demonstrated that intracellular Ca^2+^ levels in R5 are elevated with greater sleep need and that this higher intracellular Ca^2+^ was critical for the synaptic plastic changes encoding sleep drive in these neurons[36]. Thus, we assessed whether thermogenetic activation of astrocytes would lead to a similar increase in R5 intracellular Ca^2+^. To do this, we first generated a QF2 driver line broadly labeling astrocytes (*alrm-QF2*) (Figures S7A and S7B). Importantly, we found that dTrpA1 activation of astrocytes led to a substantial elevation of GCaMP signal in the R5 neurons (Figures 7H and 7I). These findings suggest that astrocytes sense substantial sleep need and convey this information, by upregulating and releasing Spz. In addition, our data support the hypothesis that Spz then activates Toll receptor in R5 neurons to generate sleep drive.

## DISCUSSION

Although astrocytes have been implicated in the homeostatic regulation of sleep[8], their specific role and the underlying mechanisms have been unresolved. Our data support a role for astrocytes as sensors of sleep need and define signaling mechanisms within these cells that mediate the integration and transmission of this information to a downstream homeostatic sleep circuit (Figure 7J). In this model, neural activity is sensed by astrocytic processes, leading to an increase in Ca^2+^ levels, which depends at least in part on specific L-type Voltage-Gated Ca^2+^ channels (VGCC)[57–60]. Interestingly, while astrocytes have been shown to exhibit hyperpolarized membrane potentials with small depolarizations[57], this particular subtype of L-type VGCC can be activated at substantially lower membrane potentials than other members of this channel family[61].

As the increased neural activity persists, Ca^2+^-mediated transcription of TyrRII is induced in astrocytes. TyrRII is relatively unstudied, but *in vitro* data suggest that it responds non-specifically to multiple monoamines[62]. Thus, its upregulation in astrocytes should sensitize these cells to signaling via monoamines, which are intimately associated with wakefulness[63]. The requirement for monoamines in this pathway may provide a logic gate for the system, imparting specificity to the signaling mechanism acting downstream of neural activity, whose semantic properties may be too broad. TyrRII itself is required for further Ca^2+^ elevations, leading to a positive-feedback loop.

Our data suggest that this amplification of astrocytic Ca^2+^ signals results in transcriptional upregulation of *spz*, the fly analog of IL-1. There is an accumulating body of evidence implicating IL-1 in sleep homeostasis in mammals[55, 64–66], and our findings demonstrating a functional role for astrocytic Spz in sleep homeostasis demonstrate that these mechanisms are conserved from invertebrates to vertebrates. Our data suggest that, under conditions of strong sleep need, Spz is released from astrocytes and transmits this information by signaling to a central sleep drive circuit (the R5 neurons) to promote homeostatic sleep “rebound.”

From a broader perspective, our model draws attention to a fundamental, yet poorly understood, aspect of sleep homeostasis—how a highly dynamic input (i.e., neural activity operating on millisecond timescale) is integrated and transformed to generate a sleep homeostatic force that functions on a significantly slower timescale. Although the precise identity of the signals embodying sleep need remain unclear, there is substantial experimental and conceptual support for the notion that neural activity increases with wakefulness[67] and is a key trigger for this process[3, 28]. Yet, the dynamic mechanisms by which this neural activity, and, by extension, sleep need is transformed to sleep drive are unknown. The homeostatic regulation of processes and behaviors involving bistable states, such as sleep vs wakefulness, requires a prominent delay between the detection of the disturbance and the generation of the response[68, 69]. In addition, the stability and switching between such bistable states can be facilitated by positive feedback loops[46, 70–72]. We propose that the transcription/translation of TyrRII, coupled with the generation of a positive-feedback loop, provide a timing delay followed by a more rapid elevation in astroglial Ca^2+^ after reaching a set threshold, thus enabling a non-linear response to the continual sampling of sleep need. The transcriptional/translation upregulation of Spz could represent an additional layer of delay.

While the sleep homeostat has been implicated in regulating sleep under both baseline conditions and following substantial sleep loss[73], the astrocytic TyrRII/Spz à R5 Toll pathway selectively regulates sleep “rebound” only after protracted sleep deprivation, as previously shown for the R5 sleep drive circuit itself[36]. Thus, this pathway likely represents a distinct process for conveying potent sleep need to generate robust sleep drive. Our findings highlight the importance and enigmatic nature of these processes for the homeostatic control of baseline sleep and suggest that similar dynamic control mechanisms may also be relevant under baseline conditions.

## ACKNOWLEDGEMENTS

We thank M. Freeman, R. Jackson, T. Littleton, C. Potter, G. Rubin, and E. Schreiter for kindly sharing reagents. We also thank the Bloomington Stock Center (supported by NIH grant P40OD018537), the Vienna *Drosophila* Stock Center (www.vdrc.at), and the TRiP at Harvard Medical School (supported by NIH grant R01GM084947) for fly stocks used in this study. We thank members of the Wu Lab for discussion. This work was supported by NINDS Center Grant P30 NS050274 for use of the Core Machine Shop and the Multiphoton Imaging Core, ERC Starting Grant 758580 (S.L.), and NIH grants K99NS101065 (M.T.), R01NS094571-03S1 (M.N.W.), R01NS100792 (M.N.W.), and R01NS079584 (M.N.W.).

## AUTHOR CONTRIBUTIONS

Conceptualization, IDB, SL, MNW; Methodology, IDB, MFK, EH, MT, MNW; Software, IDB, MFK; Investigation, IDB, MFK, E-SB, EH, KP, SL, HI, MCW, MB, MT; Writing – Original Draft, IDB, MNW; Writing – Review & Editing, IDB, MFK, E-SB, EH, KP, SL, HI, MB, MCW, MT, SL, MNW; Resources, SL, MNW; Visualization, IDB, MFK, EH, E-SB; Supervision, IDB, MT, SL, MNW; Funding Acquisition, MT, SL, MNW.

## SUPPLEMENTAL DATA

**Figure S1.**
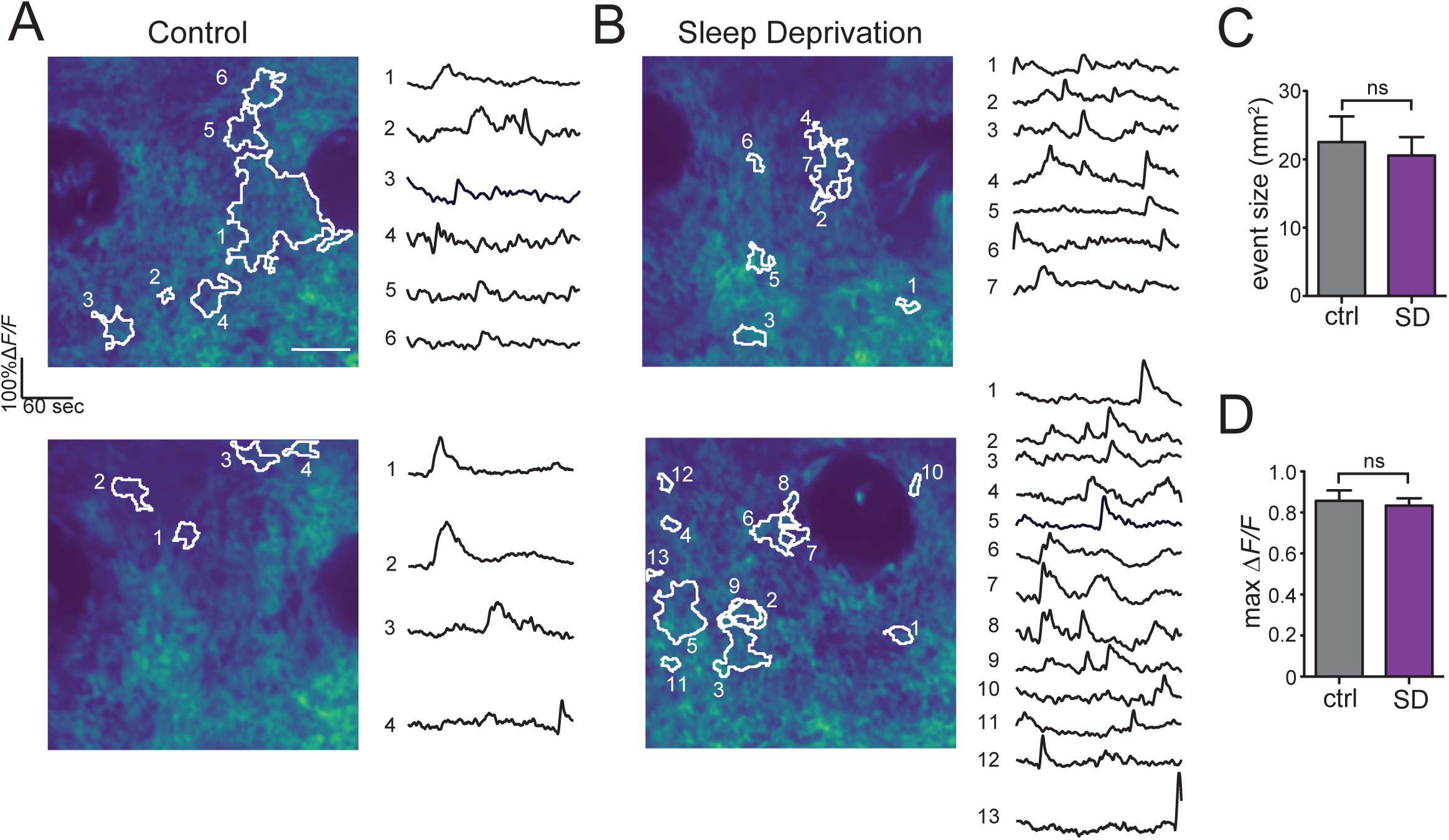
Ca^2+^ signaling in astrocytes correlates with sleep debt, Related to Figure 1. (A and B) Additional representative examples of *in vivo* GCaMP 2-photon images (left) and event traces (right) of SMP from control (A) and sleep-deprived (B) *R86E01-GAL4>UAS-myr-GCaMP6s* flies. (C and D) Event size (C), and maximum ΔF/F (D) for control (gray, n=56) and sleep-deprived (magenta, n=74) flies expressing membrane-bound GCaMP6s in astrocytes.

**Figure S2.**
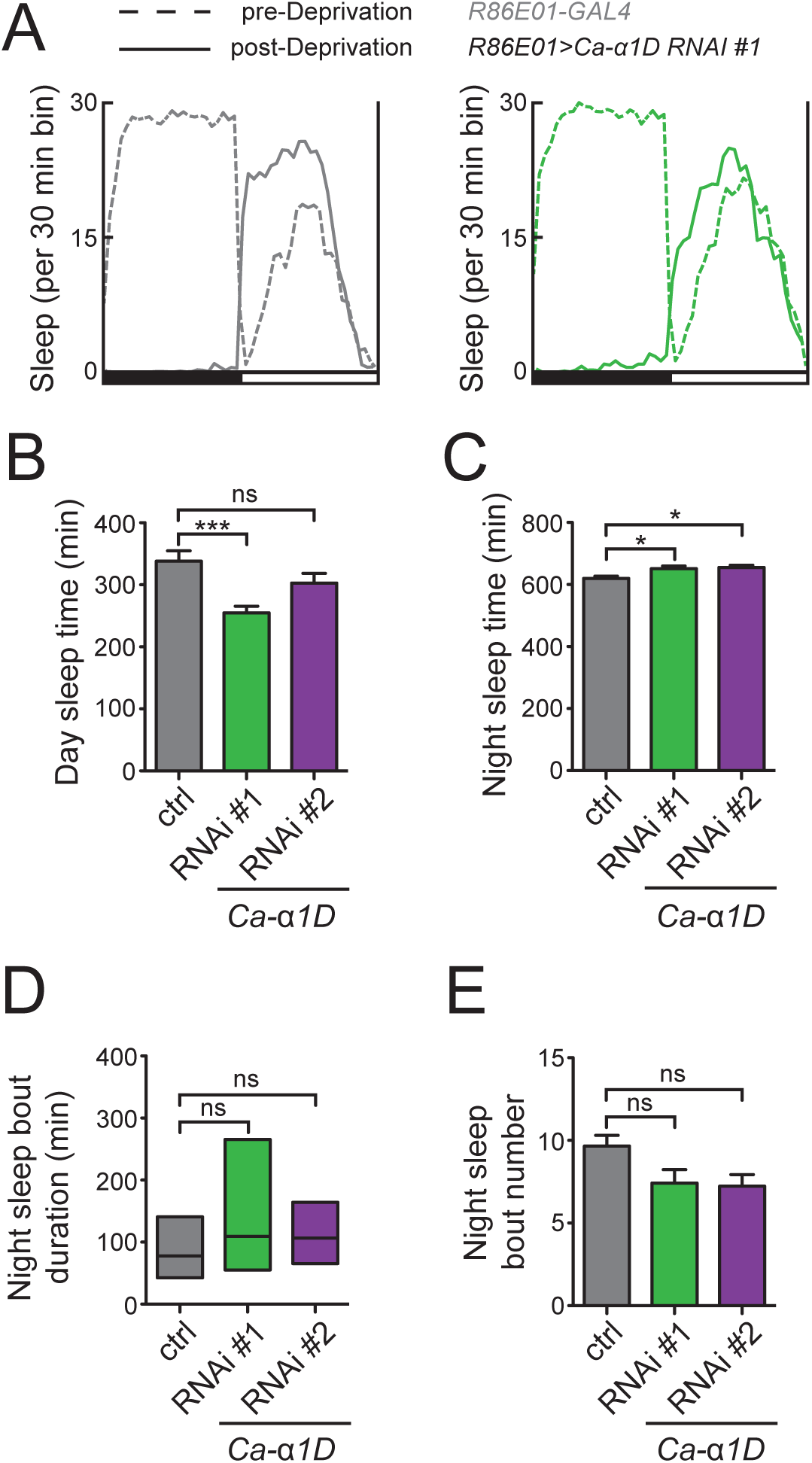
Ca^2+^ signaling in astrocytes is required for sleep homeostasis, Related to Figure 2. (A) Average 24 hr sleep profiles from baseline and post-sleep deprivation days overlaid in one plot from *R86E01-GAL4>ctrl* (gray) and *R86E01-GAL4*>*UAS-Caα1D-RNAi#1* (green) flies. White and black bars indicate 12 hr light and dark periods. (B-E) Daytime sleep time (B) nighttime sleep time (C), nighttime sleep bout duration (D), and nighttime sleep bout number (E) under baseline conditions for *R86E01-GAL4>ctrl*, *R86E01-GAL4*>*UAS-Ca-α1D-RNAi#1*, and *R86E01-GAL4*>*UAS-Ca-α1D-RNAi#2* flies. Note that the data shown in panels (A-E) are from the same animals shown in Figures 2B-2D.

**Figure S3.**
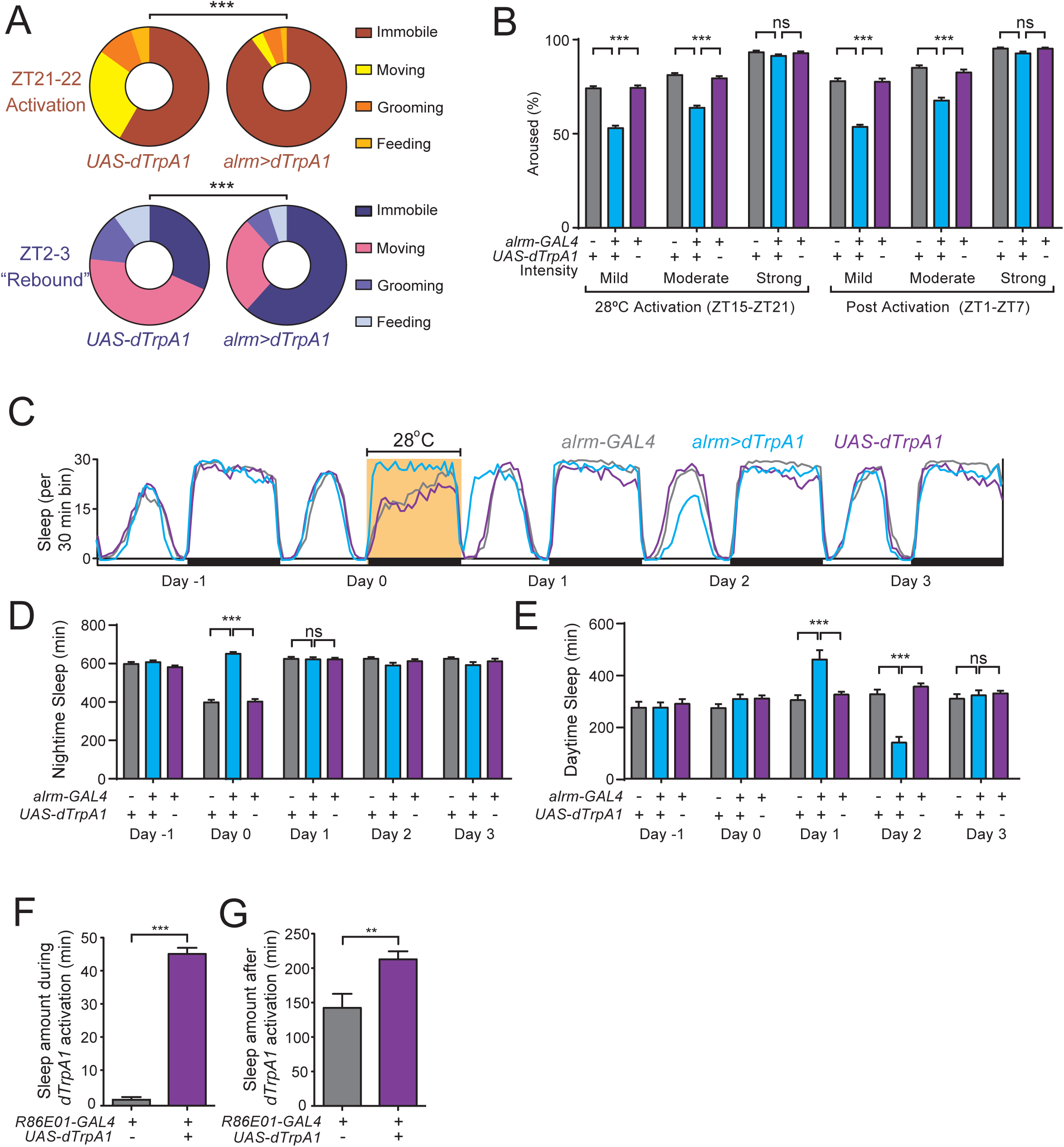
Additional behavioral characterizing sleep upon thermogenetic activation of astrocytes, Related to Figure 3. (A) Manual behavioral scoring of 1 hr of high resolution videos of *ctrl>UAS-dTrpA1* (left, n=12) and *alrm-GAL4>UAS-dTrpA1* (right, n=12) flies during thermogenetic activation of astrocytes (ZT21-22, 28°C, top) and after the temperature is returned to baseline conditions (ZT2-3, 22°C, bottom). (B) Percentage of *ctrl>UAS-dTrpA1* (n=28), *alrm-GAL4>UAS-dTrpA1* (n=28), or *alrm-GAL4* (n=29) flies aroused by a mild, moderate, or strong mechanical stimulus during thermogenetic activation of astrocytes (ZT15-21, 28°C, left) and after the temperature is returned to baseline conditions (ZT1-7, 22°C, right). (C) Sleep profile of *alrm-GAL4>ctrl* (gray), *alrm-GAL4>UAS-dTrpA1* (blue) or *ctrl>UAS-dTrpA1* (purple) flies. Highlighted period denotes 12 hr dTrpA1 activation at 28°C. (D and E) Total nighttime (D) or daytime (E) sleep amounts across experimental days for *ctrl>UAS-dTrpA1* (n=32), *alrm-GAL4>UAS-dTrpA1* (n=32), or *alrm-GAL4>ctrl* (n=31) flies shown in (C). (F and G) Sleep amount during (F) and after (G) 1 hr dTrpA1 activation from *R86E01-GAL4*>*ctrl* (n=16), and *R86E01-GAL4>UAS-dTrpA1* (n=32) flies.

**Figure S4.**
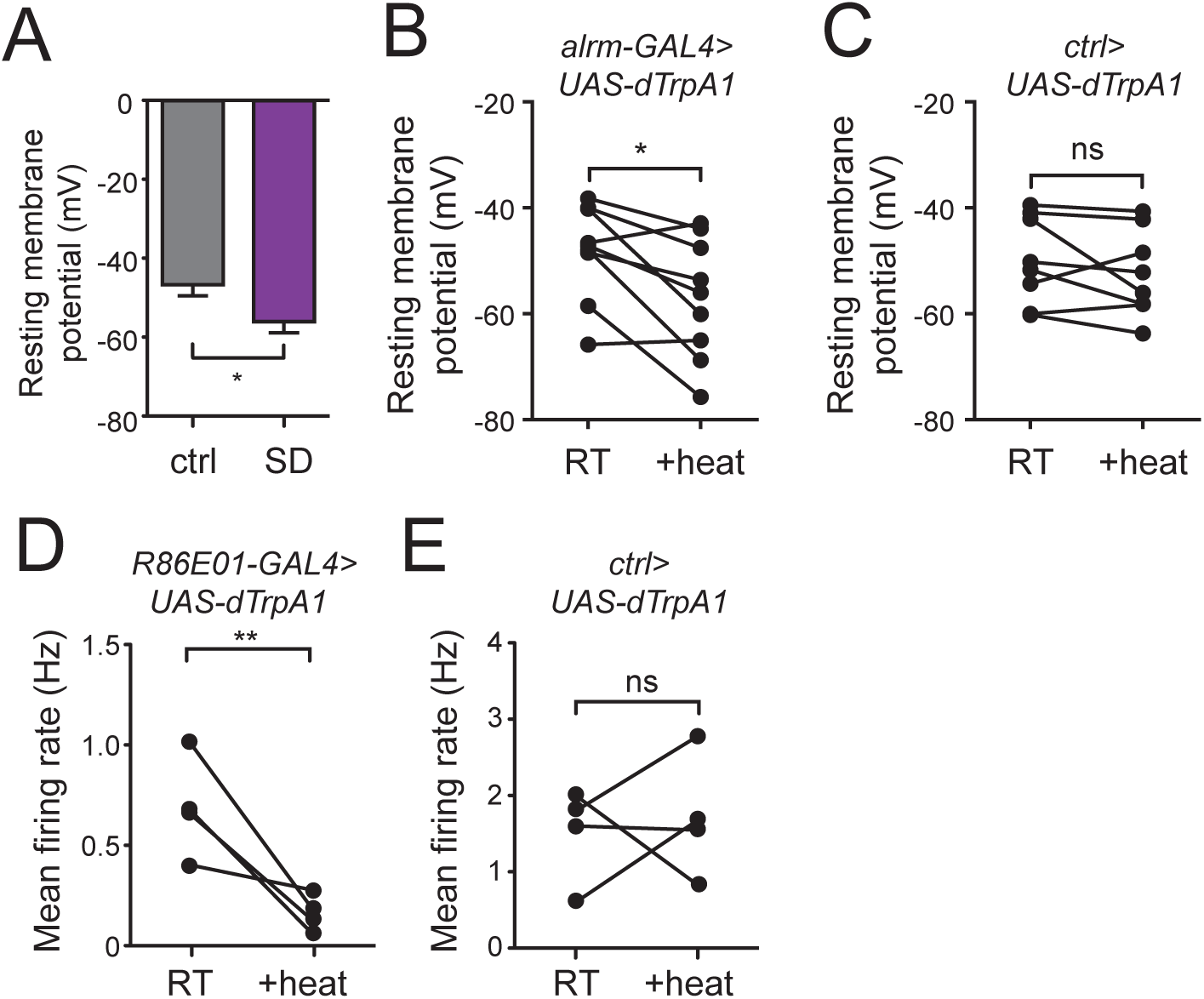
Additional electrophysiological data, Related to Figure 3. (A) Resting membrane potential of l-LNvs is significantly reduced after 12 hrs of overnight sleep deprivation. The data shown in (A) are from the same animals shown in Figures 3G and 3H. (B and C) Resting membrane potential is hyperpolarized in l-LNVs of *alrm-GAL4>UAS-dTrpA1;PDF-LexA>LexAop2-CD2::GFP* (B) but not *ctrl>UAS-dTrpA1;PDF-LexA>LexAop2-CD2::GFP* (C) control flies at 29°C (“+heat”), compared to 22°C (RT). The data shown in panels (B and C) are from the same animals shown in Figures 3I-3K. (D and E) Cell-attached recordings of l-LNvs, showing mean firing rate in *R86E01-GAL4>UAS-dTrpA1; PDF-LexA>LexAop2-CD2::GFP* flies (D) and *ctrl*>*UAS-dTrpA1;PDF-LexA>LexAop2-CD2::GFP* controls (E), before and after thermogenetic activation from room temperature (“RT”) to 28°C (“+heat”).

**Figure S5.**
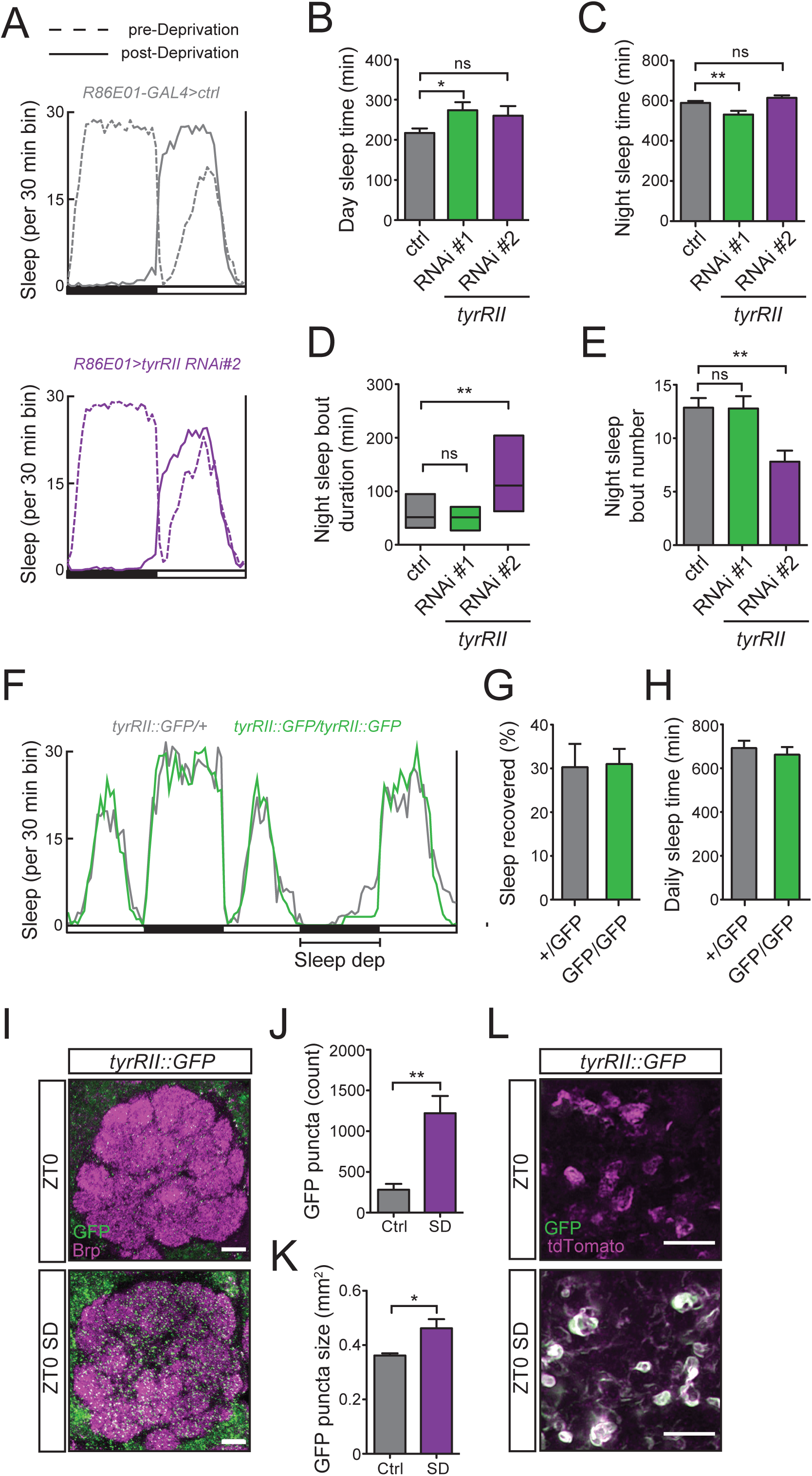
Additional behavioral and imaging data for TyrRII, Related to Figure 4 and Figure 6. (A) Average 24 hr sleep profiles from baseline and post-sleep deprivation days overlaid in one plot from *R86E01-GAL4>ctrl* (gray) and *R86E01-GAL4*>*UAS-TyrRII-RNAi#1* (green) flies. White and black bars indicate 12 hr light and dark periods. (B-E) Daytime sleep time (B), nighttime sleep time (C), nighttime sleep bout duration (D), and nighttime sleep bout number (E) under baseline conditions for *R86E01-GAL4>ctrl*, *R86E01-GAL4>UAS-TyrRII-RNAi#1,* and *R86E01-GAL4>UAS-TyrRII-RNAi#2* flies. Note that the data shown in panels (A-E) are from the same animals as shown in Figures 4D-4F. (F) Average sleep profile across baseline day and sleep deprivation from *Mi{PT-GFSTF.2}TyrRII^MI12699^* heterozygote and homozygote siblings suggests that the GFP tag does not disrupt TyrRII function. (G and H) Sleep recovered (%) (G) and daily sleep amount (H) for heterozygous (n=23) and homozygous *Mi{PT-GFSTF.2}TyrRII^MI12699^* (n=26) flies. (I) Representative images of TyrRII::GFP signal at the AL from *Mi{PT-GFSTF.2}TyrRII^MI12699^/+* flies. Whole-mount brains were immunostained with anti-GFP (green) and anti-BRP (magenta) at ZT0-1 in the presence (SD, n=5) or absence (n=5) of 12 hr sleep deprivation. (J and K) Number (J) and size (K) of TyrRII::GFP puncta in the AL at ZT1 under baseline (ctrl) or SD conditions. (L) Representative Airyscan super-resolution images of TyrII::GFP signal within the AL neuropil from *Mi{PT-GFSTF.2}TyrRII^MI12699^/+, R86E01-GAL4>UAS-mCD4::tdTomato* immunostained with anti-GFP (green) and anti-tdTomato (magenta) at ZT0-1 in the presence (SD) or absence of 12 hr SD. Scale bars denote 20 μm in (I) and 5 μm in (L).

**Figure S6.**
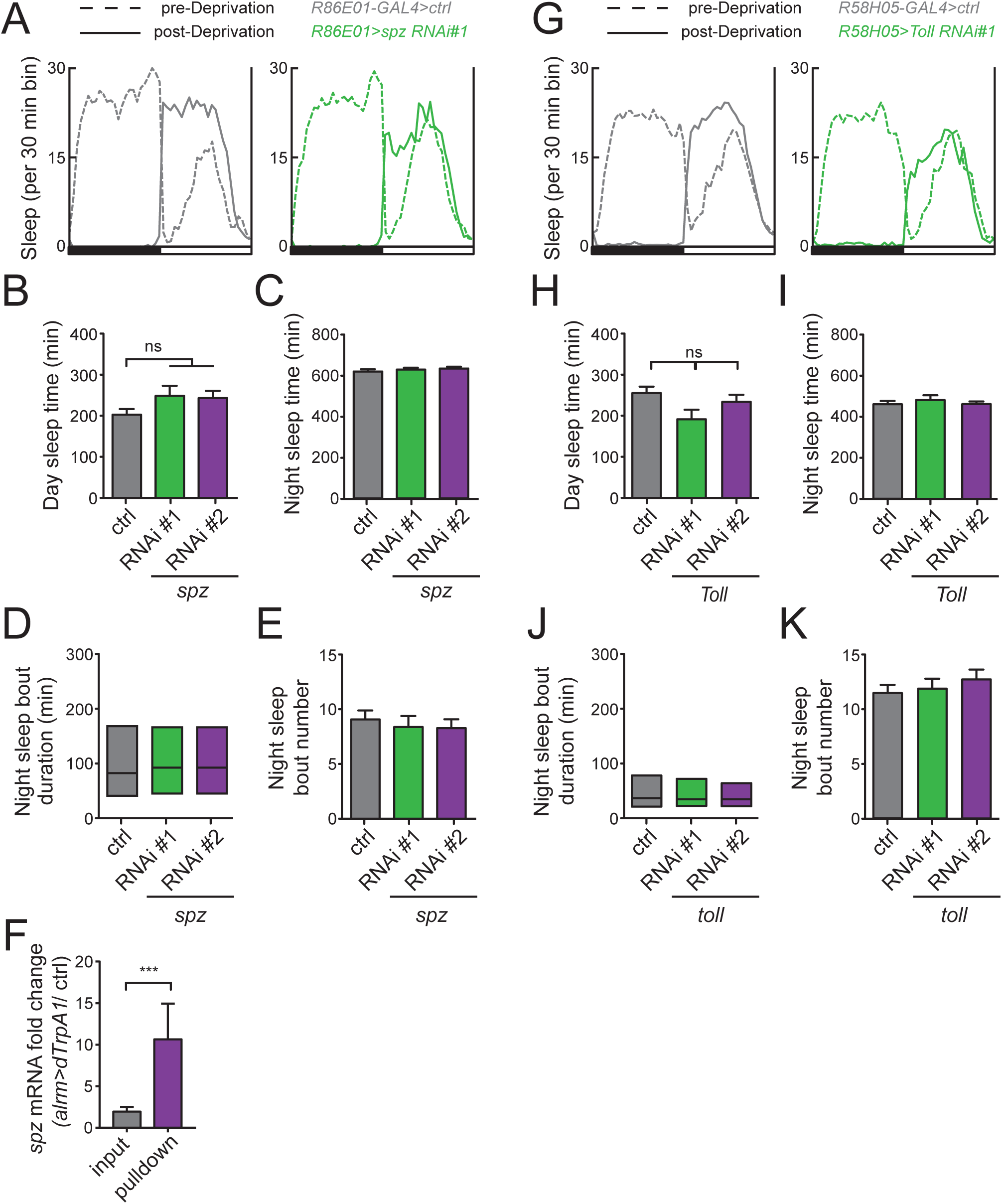
Spz in astrocytes and Toll receptor in R5 neurons are required for sleep homeostasis, but not baseline sleep, Related to Figure 7. (A) Average 24 hr sleep profiles from baseline and post-sleep deprivation days overlaid in one plot from *R86E01-GAL4>ctrl* (gray) and *R86E01-GAL4*>*UAS-spz-RNAi#1* (green) flies. White and black bars indicate 12 hr light and dark periods. (B-E) Daytime sleep time (B) nighttime sleep time (C), nighttime sleep bout duration (D), and nighttime sleep bout number (E) under baseline conditions for *R86E01-GAL4>ctrl*, *R86E01-GAL4>UAS-spz-RNAi#1,* and *R86E01-GAL4>UAS-spz-RNAi#2* flies. Note that the data in panels (A-E) are from the same animals as in Figures 7A-7C. (F) Relative fold change in *spz* mRNA level in thermogenetically activated *alrm-GAL4>UAS-dTrpA1* (n=3 replicates) vs *ctrl>UAS-dTrpA1* (n=3 replicates) flies from astrocyte-TRAP (pulldown) vs whole head (input) samples. Control *repo/nSyb* ratios for this experiment are provided in Figure 5D. (G) Average 24 hr sleep profiles from baseline and post-sleep deprivation days overlaid in one plot from *R58H05-GAL4>ctrl* (gray) and *R58H05-GAL4*>*UAS-Toll-RNAi#1* (green) flies. White and black bars indicate 12 hr light and dark periods. (H-K) Daytime sleep time (H), nighttime sleep time (I), nighttime sleep bout duration (J), and nighttime sleep bout number (K) under baseline conditions for *R58H05-GAL4>ctrl*, *R86E01-GAL4>UAS-Toll-RNAi*, and *R86E01-GAL4>UAS-Toll-miR* flies. Note that data from panels (G-K) are from the same animals as shown in Figures 7E-7G.

**Figure S7.**
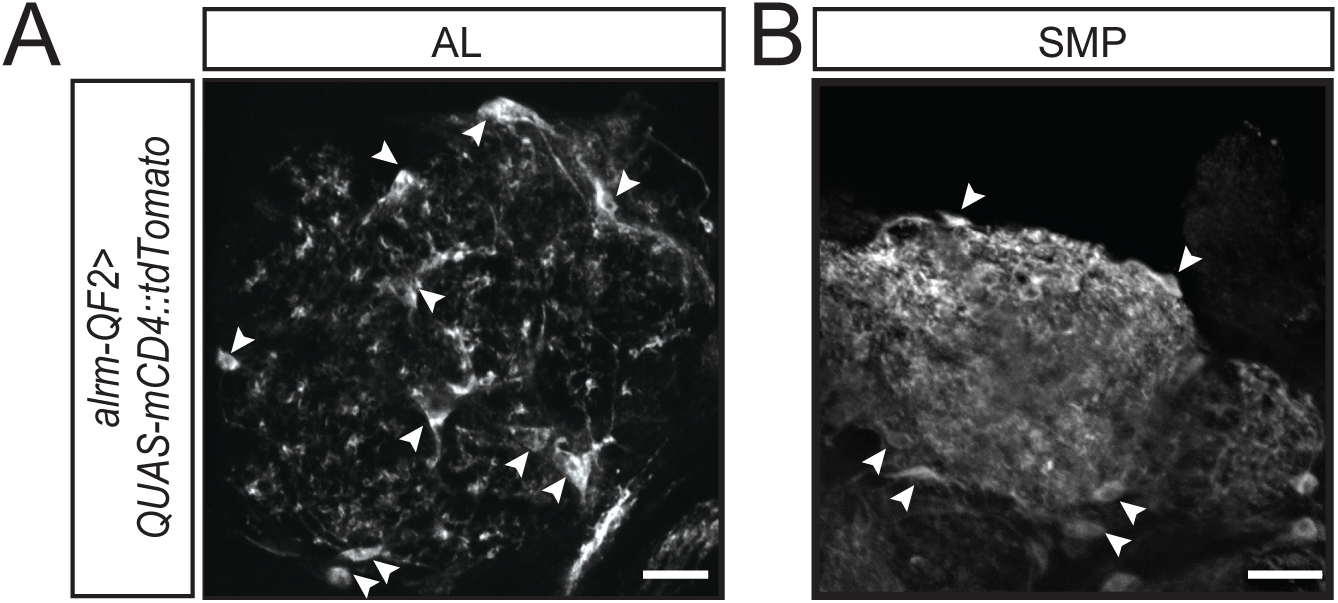
Genetic expression patterns of QF2 lines generated in this study, Related to Figure 7. (A and B) *alrm-QF2* expression recapitulates *alrm-GAL4* labeling. *alrm-QF2* labels astrocytes throughout the brain as demonstrated in the AL (A) and SMP (B) in *alrm-QF2>QUAS-mCD4::mtdTomato* flies, immunostained using anti-dsRed antibodies. White arrows identify cell bodies with distinct astrocyte-like morphology. Scale bars denote 20μm.

**Table S1.** Nomenclature, genotypes and sources for fly strains used in this study.

| Name | Source/ID# | Notes/Reference |
| --- | --- | --- |
| *alrm-GAL4* | Bloomington #67032 |  |
| *alrm-QF2 (J)* | Wu Lab | 2nd Chr random insertion |
| *GMR58H05-AD* | Wu Lab | PhiC31 mediated insertion |
| *GMR46C03-DBD* | Wu Lab | PhiC31 mediated insertion |
| *GMR72G06-GAL4* | Bloomington #39792 |  |
| *GMR86E01-GAL4* | Bloomington #45914 | Astrocyte Driver Identified by Kremer et al., 2017 *Glia* |
| *Iso31* | Bloomington #5905 |  |
| *LexAop2-CD2::GFP* | Bloomington # 66687 |  |
| *Pdf-GAL4* | Bloomington #6900 |  |
| *Pdf-LexA* | Gift from M. Rosbash |  |
| *QUAS-dTrpA1* | Gift from C. Potter |  |
| *QUAS-GCaMP6s* | Gift from C. Potter |  |
| *QUAS-mtdTomato::3xHA* | Bloomington #30005 |  |
| *Mi{PT-GFSTF.2}TyrRII<sup>MI12699-GFSTF.2</sup>* | Bloomington #65339 | Nai-Jagarwal et al., 2015 *eLife* |
| *UAS-Ca alpha 1D RNAi #1* | Bloomington #25830 |  |
| *UAS-Ca alpha 1D RNAi #2* | Bloomington #33413 |  |
| *UAS-CaMPARI2 (L398T)* | Gift from E. Schreiter | Moeyaert et al., 2018 *Nature Communication* |
| *UAS-dTrpA1<sup>attp18</sup>* | Gift from G. Rubin | Aso et al., 2014 *eLife* |
| *UAS-dTrpA1<sup>attp16</sup>* | Bloomington #26263 |  |
| *UAS-mCD8::GFP* | Bloomington # 5137 |  |
| *UAS-mCD4::tdTomato* | Bloomington #35837 |  |
| *UAS-myr-GCaMP6s* | Gift from T. Littleton |  |
| *UAS-RNAi (various)* | TRiP | Perkins et al., 2015 *Genetics* |
| *UAS-RNAi (various)* | VDRC | Dietzl et al., 2007 *Nature* |
| *UAS-Rpl10a::EGFP* | Bloomington #42683 | Gift from F. R. Jackson |
| *UAS-Spz RNAi #1* | Bloomington #34699 |  |
| *UAS-Spz RNAi #2* | VDRC #105017 |  |
| *UAS-TNT* | Bloomington #28838 |  |
| *UAS-Toll-miR (B)* | Wu Lab | 2nd Chr random insertion |
| *UAS-Toll-RNAi #1* | Bloomington #35628 |  |
| *UAS-TyrRII RNAi #1* | Bloomington #64964 |  |
| *UAS-TyrRII RNAi #2* | VDRC #110525 |  |
| *UAS-wtrw (2.23)* | Gift from M. Freeman | Ma et al., 2016 *Nature* |

## METHODS

### Fly Strains

Flies were maintained on standard food containing molasses, cornmeal, and yeast at room temperature. Mated female flies backcrossed at least 5 generations against the *iso^31^* strain or generated in the *iso^31^* background[74] were used for all experiments. See Table S1 for nomenclature, genotypes and sources for fly strains used. Unless otherwise stated, all *UAS-RNAi* lines used in this screen were obtained from the Vienna *Drosophila* Resource Center (VDRC, www.vdrc.at) or the Transgenic RNAi Project (TRiP) at Harvard Medical School.

### Nomenclature

Because two different groups of ellipsoid body (EB) rings neurons have been named “R2” neurons[75, 76], for clarity we have adopted the EB ring nomenclature outlined by Omoto et al.[77] and refer to the sleep drive neurons we previously characterized[36] as “R5” neurons.

### Molecular Biology

The *alrm-QF2* line was generated by subcloning the enhancer region into pPTQF#7-hsp70 (Addgene# 46136) using EcoRI and BamHI. The *R58H05-AD* and *R46C03-DBD* lines were generated by subcloning their respective enhancer regions into pBPp65ADZpUw (Addgene# 26234) or pBPZpGAL4DBDUw (Addgene# 26233), using Gateway cloning (Thermo Fisher). The following primers were used for PCR amplification of enhancers: *R58H05-F* 5’ - ATT ACC ATG CTG GAC CGG GTG CAA GG - 3’; *R58H05-R* 5’ - CTC ACA AGT CAT GGC CCT AAC GAG G - 3’; *alrm-F* 5’ - GAT CGA TCG CGG CCG CTA GTG GCG ATC CTT TCG CTC G - 3’; *alrm-R* 5’ - GAT CGG TAC CGA GTT AAT ATG GTG GGA ACT GC - 3’; *R46C03-F* 5’ - GAT CAA AGT TTG GGG CAA CTA CCC T - 3’; and *R46C03-R* 5’ - GGT TCC CGC AAA GTT AAT CTC CTG T - 3’. The construct for *UAS-Toll-miR* was generated as previously described[45]. Two 22mers bridging the 1st and 2nd coding exons of the neural transcript of the *Toll* gene (TCT CGA ACT AAG GGC AAA TAT C and GGC GAG GGC TAC AAC AAT AAT C) were used to create the two hairpin loops. The entire microRNA construct was synthesized *in vitro* (GeneArt) and then subcloned into pUAST using EcoRI and NotI. Transgenic lines were generated in the *iso^31^* background either through P-element mediated random insertion (*alrm-QF2* and *UAS-Toll-miR*) or site-directed PhiC31-mediated insertion into the *86Fb* (*R58H05-AD*) and *vk27* (*R46C03-DBD*) insertion sites (Rainbow Transgenics).

### Behavioral Analyses

#### Baseline sleep measurements

Sleep data collection was performed as previously described[45]. All analysis was performed using custom Matlab scripts (Mathworks) according to previously established algorithms[36]. Briefly, CO_2_-anesthetized 3-4 day old female flies were loaded into 5×65mm diameter glass tubes with 5% sucrose in agar and sealed with wax and yarn, and then allowed to recover for ∼1.5 days prior to data collection. Flies were loaded into *Drosophila* Activity Monitoring System devices (DAMS, Trikinetics), which were placed in incubators at 22°C with independent lighting control (12 hr:12 hr L:D cycles). Sleep epochs were identified based on the previously established criterion of 5 contiguous min of locomotor activity quiescence[78].

#### Thermogenetic activation

For all thermogenetic activation experiments, *UAS-dTrpA1* on the 2^nd^ chromosome was used, and activation was performed by ramping the temperature of the incubator according to the schedules described herein. Continuous temperature monitoring revealed an ∼30 min lag for temperatures to stably reach desired setpoints, which is reflected in the times used for analysis of sleep during activation for the 1 hr pulse (i.e., ZT0.5-ZT1.5).

#### Mechanical sleep deprivation

Flies were mechanically stimulated for 2-10 s/min from ZT12-ZT24 using a vortexer mounting plate and multi-tube vortexer (Trikinetics). Only data from flies with ≥90% reduction in sleep amount during deprivation, compared with baseline conditions, were included for analysis. “% Sleep Recovered” was calculated by using a sliding 30 min window to subtract baseline sleep from post-deprivation sleep binwise and then summing each with all previous bins to provide a cumulative tally of sleep time over the twelve hours post-deprivation. We then divided each 30 min bin by the total sleep lost (nighttime sleep during baseline – nighttime sleep during deprivation) and converted this ratio to a percentage.

#### Arousal threshold analysis

Flies were mechanically stimulated for 1 s/hr from ZT15-ZT21 and again from ZT1-ZT7 using a vortexer mounting plate and multi-tube vortexer (Trikinetics). The intensity of stimulus was varied using the speed adjustment knob of the vortexer from 1 (mild) to 3 (moderate) to 7 (strong) with two consecutive hourly pulses at each respective intensity, in that order. These brief arousals were performed with concomitant overnight thermogenetic activation (28°C) from ZT12-24 and a return to baseline temperatures from ZT0-12 (22°C). Flies that were inactive for 5 min before a stimulus and exhibited beam crossings within 3 min after the mechanical stimulus were identified as “aroused” and where possible the two repeats for each animal at each intensity and temperature condition were averaged. The percentage was calculated as the number of animals aroused compared to all potentially arousable animals (i.e., inactive for 5 min prior to stimulus) and each experiment was performed in triplicate.

#### RNAi screens

For the Ca^2+^ effector miniscreen, we identified genes encoding proteins that flux Ca^2+^, including ionotropic receptors, channels, transporters, and exchangers. *UAS-RNAi* lines for these genes were crossed to *R86E01-GAL4*, and the appropriate progeny (n=8) were assessed for sleep rebound phenotypes following mechanical SD from ZT12-ZT24, as described above. For our large-scale screen ∼3,200 *UAS-RNAi* lines were selected for genes that were either randomly selected or identified from previous mammalian and invertebrate astrocyte expression studies[51, 79]. These *UAS-RNAi* lines were crossed to *alrm-GAL4>UAS-dTrpA1* flies, and the appropriate progeny (n=4) were assessed for “sleep rebound” phenotypes for the 6 hrs following a 1 hr 31°C heat pulse from ZT0-1. Lines exhibiting sleep parameters ≥2.5 SD less than the mean were selected for secondary screening and further characterization.

#### Video analysis

Flies were loaded into 3D-printed Raspberry Pi-enabled recording chambers (Ethoscopes) and recorded using high-resolution video mode[80]. After two baseline days for acclimatization, the animals were recorded for 2 hrs during thermogenetic activation (ZT 20-22, 28°C) and 2 hrs after activation (ZT 2-4, 22°C). The first hour of each recording was used for manual scoring of behavior using open-source Behavioral Observation Research Interactive Software (BORIS)[81]. Behaviors were categorized into 4 possible states each minute. In cases where multiple behaviors were observed within the same minute, they were classified in this order: feeding>grooming>locomotion>immobility such that only the highest order state was labelled regardless of the presence of the other behavioral states.

### Imaging

#### Confocal Microscopy

All confocal images were acquired from cover-slipped (size 1.5) and Vectashield (Vector Labs) mounted samples using an LSM710 microscope and Zen Black image capture software (Zeiss International), except for CaMPARI and super-resolution images which were performed on an inverted LSM800 fitted with Airyscan detectors. In the latter case, Airyscan detectors were calibrated to the brightest signal from the experimental condition and then these settings were used for all samples. In all cases, pinhole aperture and slice thickness were optimized according to the software recommendations for lens NA, magnification, and reported XY resolution.

#### Ex vivo CaMPARI imaging

4-5 day old flies were loaded into locomotor tubes for sleep recordings +/- mechanical deprivation (as described above). Animals were removed between ZT0-ZT3 or ZT12-15 and quickly dissected in Adult Hemolymph-Like Saline (AHLS, 103 mM NaCl, 3 mM KCl, 1.5 mM CaCl_2_, 4 mM MgCl_2_, 26 mM NaHCO_3_, 1 mM NaH_2_PO_4_, 10 mM trehalose, 10 mM glucose, 5 mM TES Buffer, 2 mM sucrose) and then transferred to glass bottomed dishes (Pelco, 14035-20). Data were collected as 1024×1024 pixel confocal stacks targeting approximately 10 μm thick slices of the Antennal Lobes (AL) and Superior Medial Lobes (SMP) centered on astrocyte cell bodies using a 40X lens (Zeiss Plan-Apochromat 63x/1.4 Oil) prior to, and immediately after, photoconversion (PC) using 20% intensity light generated by a TTL LED (Excite) filtered with a 395/25 nm bandpass filter. Photoconversion was achieved by quickly cycling the LED with a 500ms/200ms duty cycle performed for 240 cycles (∼2.8 minutes total). Data were analyzed by first applying the “TurboReg” plugin of ImageJ (NIH) to align pre- and post-PC images and then manually drawing ROIs covering individual astrocyte cell bodies and a single neuropil region using the green channel of pre-PC images. Data analyses and generation of heat-mapped images were performed with a custom macro written for ImageJ (.ijm file available upon request) by first performing maximal intensity projection and then applying the photoconversion algorithm used by Moeyaert et al., 2018[34]. Briefly, “Fold R/G” for images represent (Red/Green)_post_ divided by (Red/Green)_pre_ at the pixel level. Average Fold R/G across each ROI was used for quantification.

#### Immunocytochemistry

Immunostaining of whole-mount brains was performed as previously described[36]. Briefly, brains were fixed in 4% PFA for ∼30 min, washed 5x in PBST (PBS + 0.3% Triton X-100), then blocked in normal goat serum, before incubation with rabbit anti-GFP (Invitrogen, 1:1000), chicken anti-GFP (Invitrogen, 1:200), rabbit anti-DsRed (Clontech, 1:1000), or mouse anti-brp (nc82, Development Studies Hybridoma Bank, 1:20), at 4°C for ∼48 hrs, followed by incubation with Alexa 488 anti-rabbit (Invitrogen, 1:1000), Alexa 488 anti-chicken (Invitrogen, 1:1000), or Alexa 568 anti-mouse (Invitrogen, 1:1000) secondary antibodies at 4°C for 2-24 hrs.

#### TyrRII::GFP quantification

Data were collected from flies expressing GFP-tagged Tyramine Receptor II as 1024×1024 pixel confocal stacks using a 40x lens (Zeiss Plan-Apochromat 40x/1.3 Oil) and were cropped to include the AL. Stacks were processed and calculated using a custom macro written for ImageJ using “Maximum Entropy Thresholding” and the “Analyze Particles” function with a 3-100 pixel cutoff to quantify GFP puncta (.ijm file available upon request).

#### In vivo astrocyte GCaMP imaging

5-6 day old flies were anesthetized on ice and placed in a hole that was etched on a stainless steel shim attached to a custom 3D-printed holder. The head capsule and thorax were glued to the shim using UV cured glue (Loctite, 3972). Legs, proboscis and antennae were immobilized using beeswax applied with a heated metal probe. The head capsule was bathed in AHLS. A small window was opened by removing the cuticle above the central brain using sharpened forceps (Dumont 5SF). Fat and other tissue were removed to gain optical access to the brain. Astrocytes expressing myristoylated GCaMP6s were imaged using 2-photon microscopy from ZT0-2 with a Zeiss LSM 710 microscope using a Ti:Sapphire Laser (@920nm, Chameleon Ultra II, Coherent). Images were acquired at 0.484 seconds a frame (∼2.1 Hz). The image window was 80 x 80 μm at 256 × 256 pixel resolution, and duration of imaging did not exceed 8 min per animal. The imaging plane was limited to the superior medial protocerebrum (SMP). Animals exhibiting spontaneous astrocytic Ca^2+^ activity were analyzed. Acquired images were first motion-corrected using a previously published method[82] with custom parameters. Motion-corrected images were processed using AQuA (Astrocyte Quantitative Analysis) which allows characterization of spatiotemporally distinct events[83]. Events were calculated based on empirically determined parameters, which were used for all images. Traces from each event were exported and filtered with a 3^rd^ order Savitzky-Golay filter over 15 frames.

#### Ellipsoid body R5 ring GCaMP imaging

For intracellular Ca^2+^ measurements of the R5 ellipsoid body ring, 5-6 day old *R58H05-QF2>QUAS-GCAMP6s, QUAS-mtdTomato-3xHA* flies bearing either *alrm-GAL4>UAS-dTrpA1 or UAS-dTrpA1* alone transgenes were examined. All animals were administered a heat stimulus (31°C) from ZT0-ZT1 and dissected and imaged from ZT3-ZT5. Brains were quickly dissected in AHLS and imaged in the same solution using an Ultima multiphoton microscope (Prairie Technologies). Excitation of both GCaMP6s and mtdTomato was achieved with 920nm light produced by a Ti:Sapphire Laser (Chameleon Ultra II, Coherent). Data were collected using a 40x water immersion lens with 2x optical detector zoom (Olympus LUMPLFLN 40XW/0.8) as a single 512×512 pixel plane over 60 s at a frequency of ∼1 Hz using Prairieview software (Prairie Technologies). The imaging plane was selected based on the completeness of the R5 ring which sits almost perpendicular to the dorsal surface of the fly brain. Data were analyzed using ImageJ by calculating the mean intensity of the R5 ring targeting ROIs (after pixel by pixel GCaMP/mtdTomato normalization) averaged over the full recording.

### Electrophysiology

3 to 9 day old flies were used in whole-cell patch clamp and cell-attached recordings. Whole-cell patch clamp recordings were generally performed as described previously[45]. For sleep deprivation experiments, flies were dissected from ZT0-2 following either 12 hrs baseline sleep or 12 hrs of mechanical SD (from ZT12-ZT24) as described above or by using the ethoscope tracking system[80], and whole-cell patch clamp recordings were performed using current-clamp mode. For astrocyte activation experiments, either whole-cell or cell-attached patch-clamp recordings were performed using current-clamp mode. Brains were dissected and recordings were performed in a *Drosophila* physiological saline solution (101 mM NaCl, 3 mM KCl, 1 mM CaCl_2_, 4 mM MgCl_2_, 1.25 mM NaH_2_PO_4_, 20.7 mM NaHCO_3_, and 5 mM glucose [pH 7.2]), pre-bubbled with 95% O_2_ and 5% CO_2_, at room temperature. If needed for removal of the glial sheath, brains were treated with 2 mg/ml protease XIV (Sigma-Aldrich) for 5–8 min at 22°C. For both whole-cell and cell-attached recordings, l-LNvs were located using GFP fluorescence or infrared-differential interference contrast (IR-DIC) optics, under a fixed-stage upright microscope (BX51WI; Olympus or SliceScope, Scientifica). Patch-pipettes (5–13 M*Ω*) were made from borosilicate glass capillary with a Flaming-Brown puller (P-1000 or P-97; Sutter Instrument) and polished with a microforge. For whole-cell recordings, the pipette was filled with internal solution containing 102 mM potassium gluconate, 0.085 mM CaCl_2_, 0.94 mM EGTA, 8.5 mM HEPES, 17 mM NaCl (pH 7.2), 4 mM Mg-ATP, and 0.5 mM Na-GTP. For cell-attached recordings, the pipette was filled with the filtered *Drosophila* physiological saline solution. For astrocyte dTrpA1 activation experiments heat stimulation was applied by perfusing solution that was preheated using a temperature controller (ThermoClamp-01, Automate Scientific, Berkeley, CA or Scientifica Temperature Controller, Scientifica) into the recording chamber. Recordings were acquired with an Axopatch 200B or Multiclamp 700B amplifier (Molecular Devices) and Digidata 1440A or Digidata 1550B interface (Molecular Devices), using pCLAMP 10 or 11. Signals were sampled at 10 or 20 kHz and low-pass filtered at 2 or 3 kHz. For each cell-attached recording, a brief electrical stimulus (15V amplitude, 10k Hz square wave, 0.05 ms duration) was applied after current-clamp recordings from each cell to verify access to the cell. The cell was discarded if this stimulus did not elicit action potentials. For the cell-attached recordings, 2 cells with a mean firing rate >9 Hz were excluded, due to concerns about cell health.

### Translating ribosomal affinity purification with quantitative polymerase chain reaction

Translating ribosomal affinity purification (TRAP) was performed as previously described using purified EGFP antibody (19C8 antibody, Memorial Sloan Kettering Cancer Center)[84] to pulldown ribosomal complexes and their associated transcripts. Briefly, ∼1,024 fly heads (per group) were collected in liquid N_2_ from 5-6 day old *R86E01-GAL4>UAS-Rpl10a::EGFP* flies at ZT0-2 under baseline conditions or immediately following 12 hrs mechanical SD from ZT12-ZT24 (described above). Following short-term storage at −80°C, heads were pulverized in homogenization buffer and then incubated with antibody-coupled magnetic beads (Dynabeads Antibody Coupling Kit, Invitrogen) to immunoprecipitate ribosomal bound mRNA species. After RNA extraction (Trizol, Invitrogen), cDNA libraries were synthesized using SuperScript III high capacity Reverse Transcription Kit (Invitrogen), and qPCR was performed using the Power SYBR Green PCR master mix (Life Technologies) using the following primers: *repo-F* 5’- GCA TCA AGA AGA AGA AGA CGA GA – 3’; *repo-R* 5’- GTT CAA AGG CAC GCT CCA - 3’; *nSyb-F* 5’- GGT CGA TGA GGT CGT GGA C – 3’; *nSyb-R* 5’- CCA GAA TTT CCT CTT GAG CTT GC -3’; *Rpl49-F* 5’- CGG ATC GAT ATG CTA AGC TGT - 3’; *Rpl49-R* 5’- CGA CGC ACT CTG TTG TCG - 3’; *spz-F* 5’- GCA ACG ATC TTC AGC CCA CG – 3’; *spz-R* 5’- TTGATCCGCTCCTTCGCACT - 3’; *TyrRII-F* 5’ – GGC TGG ATA CTG TGC GAC AT – 3’; *TyrRII-R* 5’- GTG ACG GCG AGA TAC CTG TC – 3’. A similar protocol was followed for the thermogenetic activation experiments instead using *alrm-QF2>QUAS-dTrpA1; R86E01-GAL4>UAS-Rpl10a::EGFP* or *ctrl>QUAS-dTrpA1; R86E01-GAL4>UAS-Rpl10a::EGFP* at ZT0-2 immediately following overnight (ZT12-24) heat pulse at 28°C.

### Statistical Analysis

All analyses were performed using Prism 7 (Graphpad). Normally distributed data were analyzed using parametric tests (Student’s t tests and one-way or two-way ANOVAs followed by Tukey’s post-hoc test) and plotted as bar graphs ± SEM, whereas data that violated the assumption of normality were analyzed using non-parametric tests (Kruskal-Wallis test followed by Dunn’s post-hoc test) and plotted as simplified boxplots (Median with 1^st^, and 3^rd^ Quartile boxes). For the video analysis, a Chi-squared analysis was performed to assess differences in the distribution of experimental *alrm>dTrpA1* group relative to expected values provided by the *UAS-dTrpA1*controls.

## REFERENCES

1. Blanchard, S., and Bronzino, J.D. (2012). Anatomy and physiology. In Introduction to Biomedical Engineering (Third Edition), J.D. Enderle and J.D. Bronzino, eds. (Boston: Academic Press), pp. 75–132.

2. Reichert, S., Pavón Arocas, O., and Rihel, J. (2019). The neuropeptide galanin Is required for homeostatic rebound sleep following increased neuronal activity. Neuron 104, 370–384.e375.

3. Cirelli, C., and Tononi, G. (2019). Linking the need to sleep with synaptic function. Science. 366, 189–190.

4. Huber, R., Ghilardi, M.F., Massimini, M., and Tononi, G. (2004). Local sleep and learning. Nature. 430, 78–81.

5. Rector, D.M., Schei, J.L., Van Dongen, H.P., Belenky, G., and Krueger, J.M. (2009). Physiological markers of local sleep. The European journal of neuroscience 29, 1771–1778.

6. Borbely, A.A., and Achermann, P. (1992). Concepts and models of sleep regulation: an overview. J Sleep Res 1, 63–79.

7. Garcia-Marin, V., Garcia-Lopez, P., and Freire, M. (2007). Cajal’s contributions to glia research. Trends Neurosci 30, 479–487.

8. Halassa, M.M., Florian, C., Fellin, T., Munoz, J.R., Lee, S.Y., Abel, T., Haydon, P.G., and Frank, M.G. (2009). Astrocytic modulation of sleep homeostasis and cognitive consequences of sleep loss. Neuron 61, 213–219.

9. Pelluru, D., Konadhode, R.R., Bhat, N.R., and Shiromani, P.J. (2016). Optogenetic stimulation of astrocytes in the posterior hypothalamus increases sleep at night in C57BL/6J mice. The European journal of neuroscience 43, 1298–1306.

10. Poskanzer, K.E., and Yuste, R. (2016). Astrocytes regulate cortical state switching in vivo. Proc Natl Acad Sci U S A 113, E2675–2684.

11. Clasadonte, J., Scemes, E., Wang, Z., Boison, D., and Haydon, P.G. (2017). Connexin 43-mediated astroglial metabolic networks contribute to the regulation of the sleep-wake cycle. Neuron 95, 1365–1380 e1365.

12. Dani, J.W., Chernjavsky, A., and Smith, S.J. (1992). Neuronal activity triggers calcium waves in hippocampal astrocyte networks. Neuron 8, 429–440.

13. Agulhon, C., Petravicz, J., McMullen, A.B., Sweger, E.J., Minton, S.K., Taves, S.R., Casper, K.B., Fiacco, T.A., and McCarthy, K.D. (2008). What is the role of astrocyte calcium in neurophysiology? Neuron 59, 932–946.

14. Papouin, T., Dunphy, J.M., Tolman, M., Dineley, K.T., and Haydon, P.G. (2017). Septal cholinergic neuromodulation tunes the astrocyte-dependent gating of hippocampal NMDA receptors to wakefulness. Neuron 94, 840–854 e847.

15. Durkee, C.A., and Araque, A. (2019). Diversity and specificity of astrocyte-neuron communication. Neuroscience 396, 73–78.

16. Berridge, M.J., Lipp, P., and Bootman, M.D. (2000). The versatility and universality of calcium signalling. Nat Rev Mol Cell Biol 1, 11–21.

17. Shigetomi, E., Patel, S., and Khakh, B.S. (2016). Probing the complexities of astrocyte calcium signaling. Trends Cell Biol 26, 300–312.

18. Bazargani, N., and Attwell, D. (2016). Astrocyte calcium signaling: the third wave. Nat Neurosci 19, 182–189.

19. Verkhratsky, A., Orkand, R.K., and Kettenmann, H. (1998). Glial calcium: homeostasis and signaling function. Physiol Rev 78, 99–141.

20. Fiacco, T.A., and McCarthy, K.D. (2006). Astrocyte calcium elevations: properties, propagation, and effects on brain signaling. Glia 54, 676–690.

21. Verkhratsky, A., and Nedergaard, M. (2018). Physiology of astroglia. Physiol Rev 98, 239–389.

22. Covelo, A., and Araque, A. (2018). Neuronal activity determines distinct gliotransmitter release from a single astrocyte. Elife 7, e32237.

23. Frank, M.G. (2013). Astroglial regulation of sleep homeostasis. Curr Opin Neurobiol 23, 812–818.

24. Freeman, M.R. (2015). *Drosophila* central nervous system glia. Cold Spring Harb Perspect Biol 7.

25. Vorster, A.P., and Born, J. (2015). Sleep and memory in mammals, birds and invertebrates. Neuroscience & Biobehavioral Reviews 50, 103–119.

26. Allada, R., and Siegel, J.M. (2008). Unearthing the phylogenetic roots of sleep. Curr Biol 18, R670–R679.

27. Vyazovskiy, V.V., Olcese, U., Lazimy, Y.M., Faraguna, U., Esser, S.K., Williams, J.C., Cirelli, C., and Tononi, G. (2009). Cortical firing and sleep homeostasis. Neuron 63, 865–878.

28. Ding, F., O’Donnell, J., Xu, Q., Kang, N., Goldman, N., and Nedergaard, M. (2016). Changes in the composition of brain interstitial ions control the sleep-wake cycle. Science. 352, 550–555.

29. Areal, C.C., Warby, S.C., and Mongrain, V. (2017). Sleep loss and structural plasticity. Current Opinion in Neurobiology 44, 1–7.

30. Niethard, N., and Born, J. (2019). Back to baseline: sleep recalibrates synapses. Nat. Neurosci. 22, 149–151.

31. Thomas, C.W., Guillaumin, M.C.C., McKillop, L.E., Achermann, P., and Vyazovskiy, V.V. (2019). Cortical neuronal activity determines the dynamics of local sleep homeostasis. bioRxiv, 756270.

32. Halassa, M.M., and Haydon, P.G. (2010). Integrated brain circuits: astrocytic networks modulate neuronal activity and behavior. Annu Rev Physiol 72, 335–355.

33. Losi, G., Mariotti, L., Sessolo, M., and Carmignoto, G. (2017). New tools to study astrocyte Ca(2+) signal dynamics in brain networks *in vivo*. Front Cell Neurosci 11, 134.

34. Moeyaert, B., Holt, G., Madangopal, R., Perez-Alvarez, A., Fearey, B.C., Trojanowski, N.F., Ledderose, J., Zolnik, T.A., Das, A., Patel, D., et al. (2018). Improved methods for marking active neuron populations. Nat Commun 9, 4440.

35. Donlea, J.M., Thimgan, M.S., Suzuki, Y., Gottschalk, L., and Shaw, P.J. (2011). Inducing sleep by remote control facilitates memory consolidation in *Drosophila*. Science. 332, 1571–1576.

36. Liu, S., Liu, Q., Tabuchi, M., and Wu, M.N. (2016). Sleep drive Is encoded by neural plastic changes in a dedicated circuit. Cell. 165, 1347–1360.

37. Rechtschaffen, A., Bergmann, B.M., Gilliland, M.A., and Bauer, K. (1999). Effects of method, duration, and sleep stage on rebounds from sleep deprivation in the rat. Sleep 22, 11–31.

38. Shang, Y., Donelson, Nathan C., Vecsey, Christopher G., Guo, F., Rosbash, M., and Griffith, Leslie C. (2013). Short Neuropeptide F is a sleep-promoting inhibitory modulator. Neuron 80, 171–183.

39. Cavanaugh, D.J., Vigderman, A.S., Dean, T., Garbe, D.S., and Sehgal, A. (2016). The drosophila circadian clock gates sleep through time-of-day dependent modulation of sleep-promoting neurons. Sleep 39, 345–356.

40. Shang, Y., Griffith, L.C., and Rosbash, M. (2008). Light-arousal and circadian photoreception circuits intersect at the large PDF cells of the *Drosophila* brain. Proceedings of the National Academy of Sciences 105, 19587–19594.

41. Parisky, K.M., Agosto, J., Pulver, S.R., Shang, Y., Kuklin, E., Hodge, J.J.L., Kang, K., Liu, X., Garrity, P.A., Rosbash, M., et al. (2008). PDF cells are a GABA-responsive wake-promoting component of the *Drosophila* sleep circuit. Neuron 60, 672–682.

42. Sheeba, V., Fogle, K.J., Kaneko, M., Rashid, S., Chou, Y.T., Sharma, V.K., and Holmes, T.C. (2008). Large ventral lateral neurons modulate arousal and sleep in Drosophila. Curr Biol 18, 1537–1545.

43. Cao, G., and Nitabach, M.N. (2008). Circadian control of membrane excitability in *Drosophila melanogaster* lateral ventral clock neurons. J Neurosci 28, 6493–6501.

44. Sheeba, V., Gu, H., Sharma, V.K., O’Dowd, D.K., and Holmes, T.C. (2008). Circadian- and light-dependent regulation of resting membrane potential and spontaneous action potential firing of *Drosophila* circadian pacemaker neurons. J Neurophysiol 99, 976–988.

45. Liu, S., Lamaze, A., Liu, Q., Tabuchi, M., Yang, Y., Fowler, M., Bharadwaj, R., Zhang, J., Bedont, J., Blackshaw, S., et al. (2014). WIDE AWAKE mediates the circadian timing of sleep onset. Neuron 82, 151–166.

46. Saper, C.B., Fuller, P.M., Pedersen, N.P., Lu, J., and Scammell, T.E. (2010). Sleep state switching. Neuron 68, 1023–1042.

47. Sehgal, A., and Mignot, E. (2011). Genetics of sleep and sleep disorders. Cell. 146, 194–207.

48. Weber, F., and Dan, Y. (2016). Circuit-based interrogation of sleep control. Nature. 538, 51–59.

49. Aston-Jones, G., and Bloom, F. (1981). Activity of norepinephrine-containing locus coeruleus neurons in behaving rats anticipates fluctuations in the sleep-waking cycle. J. Neurosci. 1, 876–886.

50. Weinberg, Z.Y., and Puthenveedu, M.A. (2019). Regulation of G protein-coupled receptor signaling by plasma membrane organization and endocytosis. Traffic 20, 121–129.

51. Ng, F.S., Sengupta, S., Huang, Y., Yu, A.M., You, S., Roberts, M.A., Iyer, L.K., Yang, Y., and Jackson, F.R. (2016). TRAP-seq profiling and RNAi-based genetic screens identify conserved glial genes required for adult *Drosophila* behavior. Front Mol Neurosci 9, 146.

52. Nagarkar-Jaiswal, S., DeLuca, S.Z., Lee, P.T., Lin, W.W., Pan, H., Zuo, Z., Lv, J., Spradling, A.C., and Bellen, H.J. (2015). A genetic toolkit for tagging intronic MiMIC containing genes. Elife 4, e08469.

53. Ma, Z., Stork, T., Bergles, D.E., and Freeman, M.R. (2016). Neuromodulators signal through astrocytes to alter neural circuit activity and behaviour. Nature. 539, 428–432.

54. Bazargani, N., and Attwell, D. (2017). Amines, astrocytes, and arousal. Neuron 94, 228–231.

55. Krueger, J.M., Nguyen, J.T., Dykstra-Aiello, C.J., and Taishi, P. (2019). Local sleep. Sleep Med Rev 43, 14–21.

56. Li, G., Forero, M.G., Wentzell, J.S., Durmus, I., Wolf, R., Anthoney, N.C., Parker, M., Jiang, R., Hasenauer, J., Strausfeld, N.J., et al. (2020). A Toll-receptor map underlies structural brain plasticity. eLife 9, e52743.

57. Letellier, M., Park, Y.K., Chater, T.E., Chipman, P.H., Gautam, S.G., Oshima-Takago, T., and Goda, Y. (2016). Astrocytes regulate heterogeneity of presynaptic strengths in hippocampal networks. Proceedings of the National Academy of Sciences 113, E2685–E2694.

58. Barres, B., Chun, L., and Corey, D. (1989). Calcium current in cortical astrocytes: induction by cAMP and neurotransmitters and permissive effect of serum factors. The Journal of Neuroscience 9, 3169–3175.

59. MacVicar, B.A. (1984). Voltage-dependent calcium channels in glial cells. Science. 226, 1345–1347.

60. Rungta, R.L., Bernier, L.-P., Dissing-Olesen, L., Groten, C.J., LeDue, J.M., Ko, R., Drissler, S., and MacVicar, B.A. (2016). Ca2+ transients in astrocyte fine processes occur via Ca2+ influx in the adult mouse hippocampus. Glia 64, 2093–2103.

61. Lipscombe, D., Helton, T.D., and Xu, W. (2004). L-Type calcium channels: the low down. Journal of Neurophysiology 92, 2633–2641.

62. Bayliss, A., Roselli, G., and Evans, P.D. (2013). A comparison of the signalling properties of two tyramine receptors from *Drosophila*. J Neurochem 125, 37–48.

63. O’Donnell, J., Ding, F., and Nedergaard, M. (2015). Distinct functional states of astrocytes during sleep and wakefulness: is norepinephrine the master regulator? Current Sleep Medicine Reports 1, 1–8.

64. Richter, C., Woods, I.G., and Schier, A.F. (2014). Neuropeptidergic control of sleep and wakefulness. Annu Rev Neurosci 37, 503–531.

65. Mackiewicz, M., Naidoo, N., Zimmerman, J.E., and Pack, A.I. (2008). Molecular Mechanisms of Sleep and Wakefulness. Annals of the New York Academy of Sciences 1129, 335–349.

66. Ingiosi, A.M., and Opp, M.R. (2016). Sleep and immunomodulatory responses to systemic lipopolysaccharide in mice selectively expressing interleukin-1 receptor 1 on neurons or astrocytes. Glia 64, 780–791.

67. Cajochen, C., Brunner, D.P., Krauchi, K., Graw, P., and Wirz-Justice, A. (1995). Power density in theta/alpha frequencies of the waking EEG progressively increases during sustained wakefulness. Sleep 18, 890–894.

68. Gupta, C., López, J.M., Ott, W., Josić, K., and Bennett, M.R. (2013). Transcriptional delay stabilizes bistable gene networks. Physical Review Letters 111, 058104.

69. Glass, L., Beuter, A., and Larocque, D. (1988). Time delays, oscillations, and chaos in physiological control systems. Mathematical Biosciences 90, 111–125.

70. Ferrell, J.E. (2002). Self-perpetuating states in signal transduction: positive feedback, double-negative feedback and bistability. Current Opinion in Cell Biology 14, 140–148.

71. Marucci, L., Barton, D.A.W., Cantone, I., Ricci, M.A., Cosma, M.P., Santini, S., di Bernardo, D., and di Bernardo, M. (2009). How to turn a genetic circuit into a synthetic tunable oscillator, or a bistable switch. PloS one 4, e8083–e8083.

72. Angeli, D., Ferrell, J.E., Jr., and Sontag, E.D. (2004). Detection of multistability, bifurcations, and hysteresis in a large class of biological positive-feedback systems. Proc Natl Acad Sci U S A 101, 1822–1827.

73. Borbély, A.A., Daan, S., Wirz-Justice, A., and Deboer, T. (2016). The two-process model of sleep regulation: a reappraisal. Journal of Sleep Research, n/a-n/a.

74. Ryder, E., Blows, F., Ashburner, M., Bautista-Llacer, R., Coulson, D., Drummond, J., Webster, J., Gubb, D., Gunton, N., Johnson, G., et al. (2004). The DrosDel collection: a set of P-element insertions for generating custom chromosomal aberrations in *Drosophila melanogaster*. Genetics 167, 797–813.

75. Hanesch, U., Fischbach, K.F., and Heisenberg, M. (1989). Neuronal architecture of the central complex in *Drosophila melanogaster*. Cell Tissue Res 257, 343–366.

76. Lin, C.Y., Chuang, C.C., Hua, T.E., Chen, C.C., Dickson, B.J., Greenspan, R.J., and Chiang, A.S. (2013). A comprehensive wiring diagram of the protocerebral bridge for visual information processing in the *Drosophila* brain. Cell Rep 3, 1739–1753.

77. Omoto, J.J., Keles, M.F., Nguyen, B.M., Bolanos, C., Lovick, J.K., Frye, M.A., and Hartenstein, V. (2017). Visual Input to the *Drosophila* Central Complex by Developmentally and Functionally Distinct Neuronal Populations. Curr Biol 27, 1098–1110.

78. Shaw, P.J., Cirelli, C., Greenspan, R.J., and Tononi, G. (2000). Correlates of sleep and waking in *Drosophila melanogaster*. Science. 287, 1834–1837.

79. Bellesi, M., de Vivo, L., Tononi, G., and Cirelli, C. (2015). Transcriptome profiling of sleeping, waking, and sleep deprived adult heterozygous Aldh1L1 - eGFP-L10a mice. Genom Data 6, 114–117.

80. Geissmann, Q., Garcia Rodriguez, L., Beckwith, E.J., French, A.S., Jamasb, A.R., and Gilestro, G.F. (2017). Ethoscopes: An open platform for high-throughput ethomics. PLOS Biology 15, e2003026.

81. Friard, O., and Gamba, M. (2016). BORIS: a free, versatile open-source event-logging software for video/audio coding and live observations. Methods in Ecology and Evolution 7, 1325–1330.

82. Pnevmatikakis, E.A., and Giovannucci, A. (2017). NoRMCorre: An online algorithm for piecewise rigid motion correction of calcium imaging data. J Neurosci Methods 291, 83–94.

83. Wang, Y., DelRosso, N.V., Vaidyanathan, T., Reitman, M., Cahill, M.K., Mi, X., Yu, G., and Poskanzer, K.E. (2018). An event-based paradigm for analyzing fluorescent astrocyte activity uncovers novel single-cell and population-level physiology. bioRxiv, 504217.

84. Heiman, M., Kulicke, R., Fenster, R.J., Greengard, P., and Heintz, N. (2014). Cell type-specific mRNA purification by translating ribosome affinity purification (TRAP). Nat Prot 9, 1282–1291.

